# Cognitive insights from evolutionarily new brain structures in prefrontal cortex

**DOI:** 10.1101/2020.11.07.372805

**Authors:** Willa I. Voorhies, Jacob A. Miller, Jewelia K. Yao, Silvia A. Bunge, Kevin S. Weiner

**Affiliations:** Helen Wills Neuroscience Institute, University of California, Berkeley, Berkeley CA, 94720 USA; Department of Psychology, University of California, Berkeley, Berkeley CA, 94720 USA

## Abstract

While the disproportionate expansion of lateral prefrontal cortex (LPFC) in humans compared to non-human primates is accepted, the relationship between evolutionarily new LPFC brain structures and uniquely human cognitive skills is largely unknown. Here, we tested the relationship between variability in evolutionarily new LPFC tertiary sulci and reasoning skills in a pediatric cohort. A novel data-driven approach in independent *discovery* and *replication* samples revealed that the depth of specific LPFC tertiary sulci predicts individual differences in reasoning skills beyond age. These findings support a classic, yet untested, theory linking the protracted development of tertiary sulci to late-developing cognitive processes. We suggest that deeper LPFC tertiary sulci reflect reduced short-range connections in white matter, which in turn, improve the efficiency of local neural signals underlying cognitive skills such as reasoning that are central to human cognitive development. To expedite discoveries in future neuroanatomical-behavioural studies, we share sulcal definitions with the field.

## Introduction

A fundamental question in cognitive neuroscience is how the structure of the brain supports complex cognition. While much progress has been made in answering this question, especially in animal models, human brains differ in both their micro- and macrostructural properties from widely used animals in neuroscience research such as mice, marmosets, and macaques^1^. These cross-species differences are especially pronounced in association cortices such as lateral prefrontal cortex (LPFC). LPFC is a late-developing cortical expanse that is enlarged in humans compared to non-human primates^2^ and is critical for cognitive control, executive function, reasoning, and goal-directed behavior^3–6^. Yet there is still much progress to be made in understanding how the development of evolutionarily new brain structures in the expanded human LPFC support the development of complex, largely human, cognitive skills achieved by neural circuits within LPFC.

Of all the cognitive skills and anatomical features to focus on, we investigate the relationship between relational reasoning and macro-anatomical structures in human cortex known as tertiary sulci. Sulci are commonly classified as primary, secondary, or tertiary based on their time of emergence in gestation^7–22^. Tertiary sulci are the last to emerge *in utero*, and subsequently are often the shallowest and smallest class of cortical folds^7–22^. They are largely overlooked due to methodological difficulties in their identification (which we expand on further below)^11,19,23^. Due to these difficulties, very little is known regarding the role of tertiary sulci in human cognition, despite the fact that many tertiary sulci are evolutionarily new structures that are uniquely human^14^. Here, we test whether tertiary sulci in LPFC are behaviorally significant: that is, whether they relate to higher cognitive functioning. We focus here on reasoning – or, more specifically, relational reasoning, which is the ability to extract common features across objects and conceptualize them in terms of their relation to each other^6,24^. Humans consistently outperform other species in tests of relational reasoning^6,25^, which relies on a distributed network involving LPFC that has expanded through primate evolution and that develops slowly over childhood and adolescence^6,26–31^. In fact, LPFC is considered critical to reasoning^6,24,30,32–35^ and developmental improvements in reasoning are correlated with, and predicted by, structural and functional connectivity between LPFC and lateral parietal cortex^24,25,30,36^.

As both reasoning skills and the LPFC exhibit protracted developmental trajectories in childhood, they serve as ideal targets to test a classic, yet largely unconsidered theory. Specifically, Sanides (1964)^9^ proposed that morphological changes in tertiary sulci would likely be associated with the slow development of higher-order thinking and cognitive skills^9,22^. Fitting these criteria, relational reasoning skills continue to develop throughout childhood, while tertiary sulci emerge late in gestation and continue to develop after birth for a still undetermined period of time^9,11,14,15,20,37–39^. A relationship between relational reasoning and tertiary sulcal morphology would build on previous findings relating the development of relational reasoning to changes in LPFC cortical thickness and structural connectivity^40,41^. Furthermore, relational reasoning supports complex problem solving and scaffolds the acquisition of additional cognitive skills in children^42,43^. Thus, exploring if or how tertiary sulci contribute to the development of this cognitive skill may not only provide insight to a classic theory, but also advance understanding of the anatomical features underlying variability in the development of a wide range of other cognitive skills.

While recent studies suggest a link between the morphology of tertiary sulci in association cortices and cognitive functions,^11,19,39,44,45,46^, no study to date (to our knowledge) has tested the role of tertiary LPFC sulci in cognitive development. This gap likely persists for three key reasons. First, previous studies examining individual differences in the development of reasoning and anatomical variability in human LPFC^24^ implemented analyses that were averaged across individuals on standard neuroanatomical templates, which obscure tertiary sulci in LPFC^19^ (Supplementary Fig. 2). Therefore, to precisely characterize the relationship between tertiary sulcal morphology in LPFC and reasoning performance, it is necessary to consider cortical anatomy at the level of the individual. Second, the shallowness of tertiary sulci makes them hard to reliably identify in post-mortem tissue—typically considered the gold standard for n e u r o anatomical analyses—because they are easily confused with shallow indentations produced by veins and arteries on the outer surface of the cerebrum^11^. Third, the patterning of LPFC tertiary sulci has remained contentious, and even detailed classic studies of LPFC sulcal patterning^16,18,21,47–51^ have left tertiary LPFC sulci undefined or conflated with surrounding structures^45^ (Online Methods). For example, the sulci consistent with the location of the *pmfs* were considered as the posterior end of the *intermediate frontal sulcus (imfs)* by Ono and colleagues^49^. However, present quantifications in adults^19^ show that this sulcus is distinct from the *imfs*. We also find that we can apply the modern atlas to define the *pmfs* in these classic images (see Appendix in Miller et al., 2021b^45^ for an historical analysis of tertiary sulci in the middle frontal gyrus). The contention in historical definitions of tertiary sulci means that neuroanatomical atlases and neuroimaging software packages largely exclude tertiary sulci. In turn, tertiary sulci in LPFC have been excluded from most developmental cognitive neuroscience studies until the present study. Fourth, a common misconception is that macro-anatomical structures such as sulci and gyri are functionally and cognitively relevant in primary, but not association, cortices. For example, the calcarine sulcus predicts the location of the primary visual cortex^52^ and a “knob” in the pre-central gyrus predicts the motor hand area in the primary motor cortex^53,54^. Nevertheless, there is increasing evidence that some tertiary sulci are functionally relevant in association cortices such as ventral temporal cortex^11^ (VTC), medial PFC^39,55^, and LPFC^19^ in adults, as well as behaviorally and clinically meaningful in medial PFC^46,56,57,58^. Despite this mounting evidence that tertiary sulci are functionally and behaviorally relevant in association cortices within adults, it is largely unknown whether morphological features of tertiary sulci will predict individual differences in behavior and cognition in a developmental cohort.

To address this gap in knowledge, we characterized LPFC tertiary sulci for the first time in a developmental sample. We studied a broad age range—children and adolescents between 6 and 18 years old—as we sought to leverage the neuroanatomical and cognitive variability intrinsically present in the sample to explore whether variability in tertiary sulcal morphology predicts individual and developmental differences in relational reasoning. As sulcal depth is a characteristic feature of tertiary sulci, which are shallower than primary and secondary sulci^1,9,11,14,15,20,37,38,57^, we hypothesized a relationship between the depth of tertiary sulci and reasoning skills.

To characterize this relationship, we developed a novel pipeline that combines the most recent anatomical definition of LPFC tertiary sulci^20^ with data-driven analyses to model sulcal morphological features and reasoning performance. Our approach addresses four main questions: 1) Can LPFC tertiary sulci be identified reliably - and if so, are they smaller, shallower, and more variable compared to primary LPFC sulci as in adults?^19,20^ 2) Is there a relationship between the depth of LPFC tertiary sulci and reasoning performance across individuals? 3) If so, can we construct a neuroanatomical-behavioral model to predict an individual’s reasoning score from tertiary sulcal depth and age in an independent sample? 4) If successful, does this neuroanatomical-behavioral model generalize to other sulcal features or cognitive tasks? Answering these questions offers the first link between tertiary LPFC sulcal morphology and reasoning, as well as provides novel cognitive insights from evolutionary new brain structures in LPFC.

## Results

### Tertiary sulci are consistently identifiable in the LPFC of 6-18 year-olds

Our sample consisted of 61 typically developing children and adolescents ages 6-18 years old. Participants were randomly assigned to *Discovery* (N = 33) and *Replication* (N = 28) samples with comparable age distributions (*Discovery*: mean(sd) = 12.0 (3.70); *Replication*: mean(sd) = 12.32 (3.53); *p* = 0.81). For each subject, we generated cortical surface reconstructions in FreeSurfer^34–36^ from high-resolution T1-weighted anatomical scans. As current automated methods do not define LPFC tertiary sulci and often include gyral components in sulcal definitions (Supplementary Fig. 2), all sulci were manually defined on the native cortical surface for each subject according to the most recent and comprehensive atlas of LPFC sulcal definitions^14^ (Fig. 2). LPFC sulci were classified as primary, secondary, or tertiary based on previous studies documenting the temporal emergence of sulci in gestation^10,15–19,21,45^. While the most modern sulcal parcellation was not included in these classic studies^10,15–19,21^, it is generally accepted that anterior middle frontal LPFC sulci emerge within the gestational window for primary sulci. Meanwhile, posterior LPFC middle frontal sulci emerge late in gestation^15,16,37^. Consequently, we designate posterior middle frontal sulci as tertiary, and all surrounding sulci as primary (see Fig. 2a for all classifications.) We describe the criteria for classification and the correspondence between historical and contemporary sulcal definitions in more detail in the *Online Methods*.

**Fig. 1.**
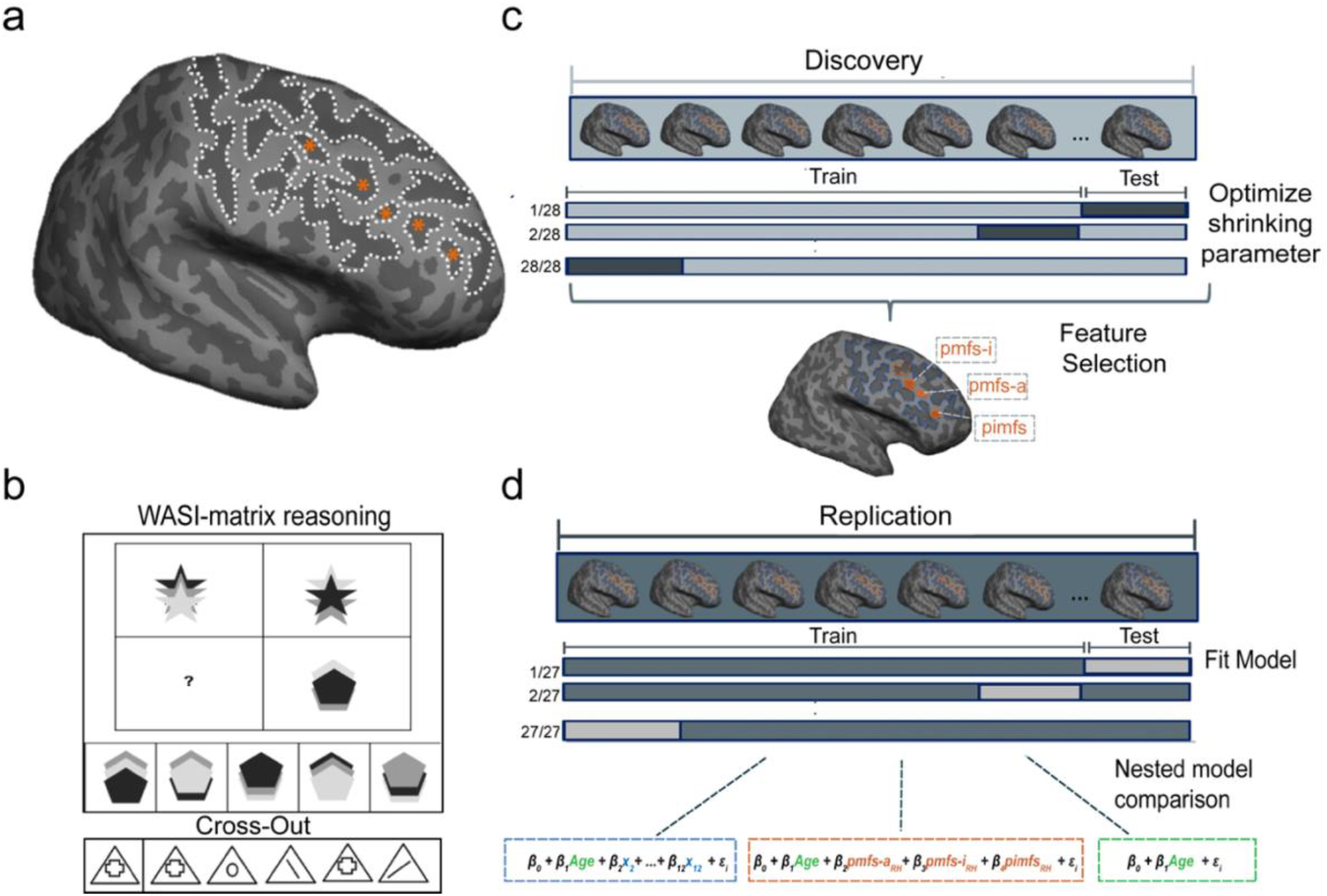
A novel, data-driven analysis pipeline with Discovery and Replication samples that models the relationship between LPFC sulcal morphological features and reasoning performance. **a.** An inflated cortical surface reconstruction of a right hemisphere from one example subject. Dotted white outlines show manually labeled sulci. Asterisks indicate the frequently omitted or misclassified tertiary sulci (Supplementary Fig. 1 for all participants). **b.** *Top:* Example from the standardized test used to assess relational reasoning in this study (WISC-IV, Matrix Reasoning task). In this task, participants are instructed to complete the matrix so that the relation between the two bottom shapes mirrors the relation between the two top shapes. In this example, option 4 completes the pattern. *Bottom:* Example from the processing speed task (WJ-R, Cross-Out test), which serves as a behavioral control. In this task, participants are instructed to cross out all objects that match the object on the left as quickly as possible. **c.** *Feature selection – Discovery sample*. A LASSO regression was performed in the *Discovery* sample to determine which sulci, if any, were associated with Matrix Reasoning performance. The model parameters were fit iteratively using a leave-one-out cross-validation procedure (Online Methods). **d.** *Model evaluation – Replication sample*. The sulci selected from the LASSO regression (orange; *pmfs-i_RH_, pmfs-a_RH_*, and *pimfs_RH_*) were included along with age in a model to predict task performance in the *Replication* sample. In order to assess the unique contribution of the selected sulci to task performance, this model (*orange*) was compared to two nested alternate models: (1) age alone (*green*) and (2) age in addition to all 12 LPFC sulci (*blue*). All models were fit with a leave-one-out cross-validation procedure.

**Fig. 2.**
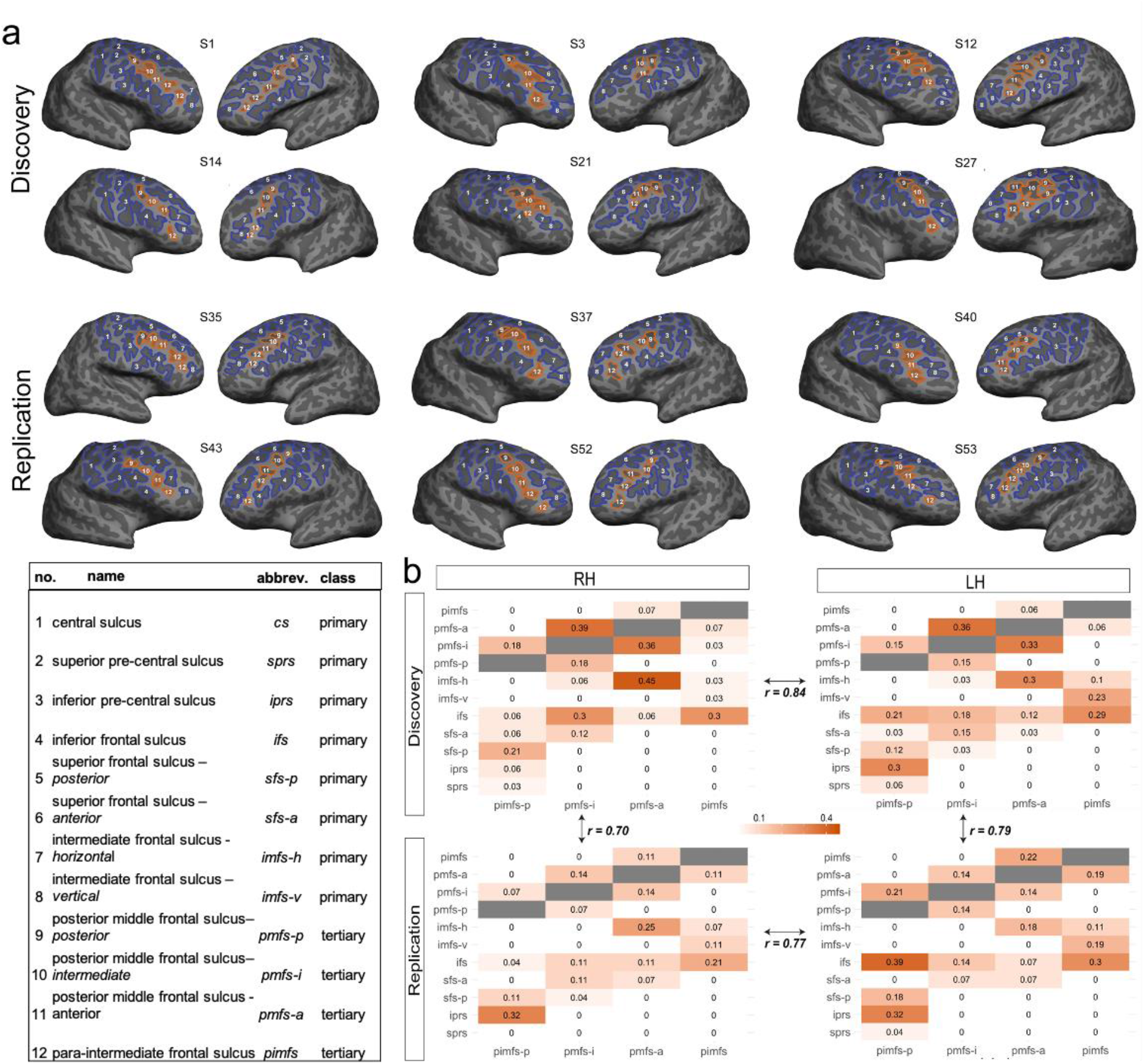
LPFC tertiary sulci are identifiable and show comparable patterning across hemispheres and samples in a pediatric cohort. **a.** LPFC sulcal definitions on inflated cortical surface reconstructions from six example participants in the *Discovery* sample (*top*) and the *Replication* sample (*bottom*). Sulci were identified based on the most recent neuroanatomical atlas to consider tertiary sulci^20^. Primary sulci (1-8) are in blue, while tertiary sulci (9-12) are in orange. The three tertiary sulci (*pmfs-i_RH_* (*10*), *pmfs-a_RH_* (*11*), and *pimfs_RH_* (*12*)) identified by our model-based approach with cross-validation (Fig. 4) are filled in. The distinction among primary, secondary, and tertiary sulci is based on classic and recent studies examining the timepoints when sulci emerge in gestation (Online Methods). Based on these studies, the sulci considered in the present study are either primary or tertiary. **b**. Rates of intersection with surrounding sulci were quantified for each tertiary sulcus in order to identify common sulcal patterns. For each tertiary sulcus (*pmfs-p* (*9*), *pmfs-i* (*10*), *pmfs-a* (*11*), and *pimfs* (*12*)), we report the proportion of intersection (frequency of occurrence/total number of observations) with each LPFC sulcus. Calculating the correlation between matrices shows that sulcal patterning is comparable (all *r*s > .70; all ps <0.0001) between hemispheres and samples.

We focused our analyses on the region commonly referred to as the dorsal LPFC, which is bounded posteriorly by the central sulcus (*cs*), anteriorly by the horizontal (*imfs-h*) and ventral (*imfs-v*) components of the intermediate frontal sulcus, superiorly by the two components of the superior frontal sulcus (*sfs-p* and *sfs-a*), and inferiorly by the inferior frontal sulcus (*ifs*). Throughout the paper, we refer to this region as the LPFC (Fig. 2a). Studies in adults report as many as five tertiary sulci within these anatomical boundaries^20^: the three components of the posterior middle frontal sulcus (posterior: *pmfs-p*; intermediate: *pmfs-i*; anterior: *pmfs-a*) and the two components of the para-intermediate frontal sulcus (ventral: *pimfs-v*; dorsal: *pimfs-d*). We defined sulci on the inflated and pial cortical surfaces of each hemisphere for each participant (Online Methods). We emphasize that 1,320 manual labels were created in total to examine the relationship between LPFC sulcal depth and reasoning performance (Supplementary Fig. 1 for sulcal definitions in all 122 hemispheres included in both samples). Sulcal definitions and all subsequent analyses are conducted separately for the *Discovery* and *Replication* samples, in order to assess the reliability and generalizability of our findings.

#### Discovery sample

All primary sulci—the central sulcus (*cs*), the superior (*sprs*) and inferior (*iprs*) portions of the precentral sulcus; *sfs-p*; *sfs-a*; *ifs*; *imfs-h*; *imfs-v*)—were identifiable in both hemispheres of each individual subject. For the first time, we demonstrate that tertiary sulci in LPFC are consistently identifiable within the hemispheres of pediatric participants as young as 6 years old (Fig. 2a). The three components of the posterior middle frontal sulcus (*pmfs-p*; *pmfs-i*; *pmfs-a*) were identifiable in all participants in every hemisphere. However, the most anterior LPFC tertiary sulcus, the *para-intermediate frontal sulcus* (*pimfs*), was consistently variable across individuals (Supplementary Table 1). Specifically, while almost all participants had at least one identifiable component of the *pimfs* (right hemisphere: 30/33; left hemisphere: 31/33), we were only able to identify both dorsal and ventral *pimfs* components in 42.42% of all participants (right hemisphere: 12/33; left hemisphere: 16/33). We further quantify this variability in tertiary sulci by examining the prevalence of sulcal types, based on their rate of intersection with neighboring sulci (Online methods; Fig. 2b). We find that sulcal patterning is very similar across hemispheres, with comparable rates of intersecting and independent sulci (r = 0.84, p<0.0001).

#### Replication sample

Consistent with the *Discovery* sample, all primary sulci (numbered 1-8 in Fig. 3b) could be identified in both hemispheres of each individual subject. In terms of tertiary sulci, the *pmfs-p, pmfs-i*, and *pmfs-a* (numbered 9-11 in Fig. 3b) were also identifiable in each hemisphere of every individual. Once again, the *pimfs* was the most variable across individuals (Supplementary Fig. 1b; Supplementary Table 1). We were able to identify at least one *pimfs* component in almost every subject (right hemisphere: 28/28; left hemisphere: 27/28). Both the dorsal and ventral *pimfs* components were identifiable in 76.8% of hemispheres (right hemisphere: 19/28 participants; left hemisphere: 24/28; Supplementary Table 1). In each hemisphere, the rates and types of intersecting sulci were highly similar to those observed in the *Discovery* sample (right hemisphere: r = 0.70, left hemisphere: r = 0.79, p<0.0001) and these were also consistent between hemispheres in this sample (r = 0.77; Figure 2b).

**Fig. 3.**
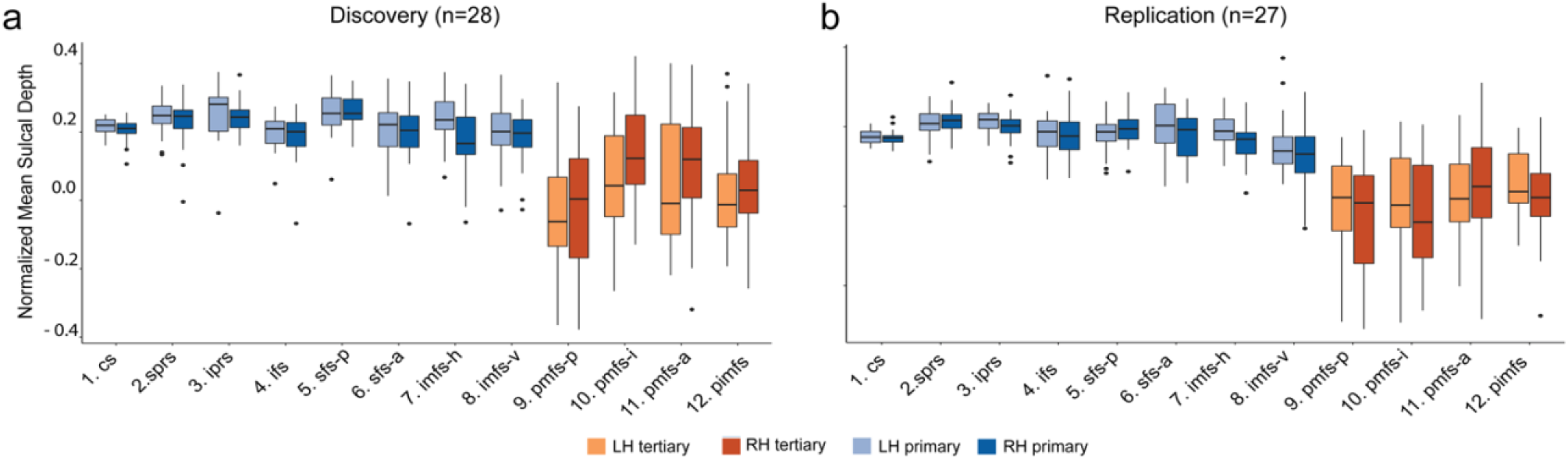
LPFC tertiary sulci in 6-18 year-olds are more shallow and more variable than primary sulci. **a.** Normalized sulcal depth for each of the 12 LPFC sulci in the *Discovery* samples. Tertiary sulci *(orange)* were shallower and more variable than primary sulci *(blue)* in both samples. **b.** Same as in ***a***. but for the *Replication* sample.

In sum, we could identify LPFC tertiary sulci in both *Discovery* and *Replication* samples and found that the sulcal patterning was comparable – and highly correlated – between each sample. However, we could not identify both dorsal and ventral *pimfs* components in each hemisphere. Thus, our inclusion criteria for all subsequent analyses was to include participants who had at least one *pimfs* component in each hemisphere (*Discovery*: 28/33, *Replication*: 27/28), which assures that all repeated measures statistics are balanced for effects of sulcus and hemisphere.

### LPFC tertiary sulci are shallower and more variable than primary sulci in children

Classic anatomical studies report a high correspondence between sulcal classification and depth ^9,14,15,37,38^, and recent *in-vivo* studies in adults show that LPFC tertiary sulci are in fact significantly shallower and more variable than primary sulci in adults^19^. However, this correspondence has not been established for LPFC sulci in children. Thus, we next sought to compare the depth and variability of LPFC tertiary and primary sulci in children. Sulcal depth was normalized to the maximum depth value within each individual hemisphere in order to account for differences in brain size across individuals and hemispheres (Online Methods). From these normalized measures, we conducted a 2-way repeated-measures analysis of variance (rm-ANOVA) to statistically test for differences between sulcal type (*tertiary vs. primary*) and hemisphere (*left vs. right*) in both *Discovery* and *Replication* samples.

#### Discovery sample

Consistent with findings in adults^19^, we observed a main-effect of sulcal type (F(1,27)= 95.63, *p*<10^-3^, *η*^2^_G_ = 0.35) in which tertiary sulci were significantly more shallow than primary sulci (Mean(sd)_Tertiary_= 0.04(0.17); Mean(sd)_Primary_= 0.23(0.07)). We also observed an interaction between sulcal type and hemisphere (F(1,27) = 5.67, *p*<0.02, *η*^2^_G_= 0.01) in which tertiary sulci were significantly deeper in the right hemisphere than in the left hemisphere (Mean(sd)_RH_ = 0.06(0.17); Mean(sd)_LH_= 0.02(0.1)). In contrast, the depth of primary sulci did not differ between hemispheres (Mean(sd)_RH_= 0.21(0.07); Mean(sd)_LH_ = 0.23(0.07)); Fig. 3a). To explore the morphological variability between sulcal types, we repeated the same analysis replacing mean sulcal depth with the standard deviation of sulcal depth. This analysis quantitatively supports that tertiary sulci are more variable than primary sulci (F(1,27)= 162.4, *p*<10^-3^, *η*^2^_G_ = 0.43), with no differences between hemispheres (*p* = 0.3).

#### Replication sample

We observed the same main effect of sulcal type in the *Replication* sample. Tertiary sulci were more shallow than primary sulci (F(1,26) = 136.5, *p*<10^-3^, *η*^2^_G_ = 0.46; Mean(sd)_Tertiary_= 0.02(0.16); Mean(sd)_Primary_ = 0.23(0.07)). We did not observe an interaction with hemisphere in this sample (F(1,26) = 0.26, *p* = 0.62); Fig. 3b). Once again, an rm-ANOVA of the standard deviation of sulcal depth revealed that tertiary sulci were more variable than primary sulci across hemispheres (F(1,26) = 170.4, *p*<10^-3^, *η*^2^_G_ = 0.47).

Additionally, while age was correlated with reasoning performance in both *Discovery* (r = 0.58, *p*<10^-3^) and *Replication* samples (r = 0.73, *p*<10^-3^), there was an inconsistent relationship between sulcal depth and age in either sample (Supplemental Fig. 4). Thus, we next implemented a two-pronged, model-based approach to test if including sulcal depth predicted reasoning skills above and beyond age.

### A model-based approach with nested cross-validation reveals that including the depth of three LPFC tertiary sulci predicts individual variability in reasoning skills above and beyond age alone

To examine the relationship between LPFC sulcal depth and reasoning skills, we implemented a data-driven pipeline with an emphasis on producing reliable and generalizable results. Based on current gold-standard recommendations^59^, we follow a four-pronged analytic approach to assess and improve the generalizability of our results and each stage of analysis (Online Methods). First, we implemented a feature selection technique in the *Discovery* sample (Fig. 1c) to determine if the depths of any LPFC sulci are associated with reasoning performance (to remind the reader, we use depth in the model because this is the main morphological difference between tertiary from primary sulci). To do so, we submitted sulcal depth values for all 12 LPFC sulci in the *Discovery* sample to a LASSO regression model, which provides an automated method for feature selection by shrinking model coefficients and removing sulci with very low coefficients from the model (Fig. 1c; Online Methods). This approach allowed us to determine, in a data-driven manner, which sulci are the strongest predictors of reasoning performance. Additionally, this technique guards against overfitting and increases the likelihood that a model will generalize to other datasets, by providing a sparse solution that reduces coefficient values and decreases variance in the model without increasing bias^59,60^. Also, although we observe a gender imbalance in our samples, gender was not associated with sulcal depth (*p* = 0.27), or Matrix Reasoning (*p* = 0.51); therefore, we do not consider gender further in our models.

To determine the value of the shrinking parameter (*α*)^60^, we iteratively fit the model with a range of *α*-values using cross-validation. By convention^41^, we selected the *α* that minimized the cross-validated Mean Squared Error (MSE_CV_; Fig. 4a). Although both tertiary and primary sulci were initially included as predictors, after implementing the LASSO regression, only three tertiary sulci (*pmfs-i, pmfs-a*, and *pimfs*) in the right hemisphere were found to be associated with reasoning performance (MSE_CV_=21.84, *α* = 0.1; *β*_pmfs-i_= 4.50, *β*_pmfs-a_=1.78, *β*_pimfs_= 11.88; Fig. 4).

**Fig. 4.**
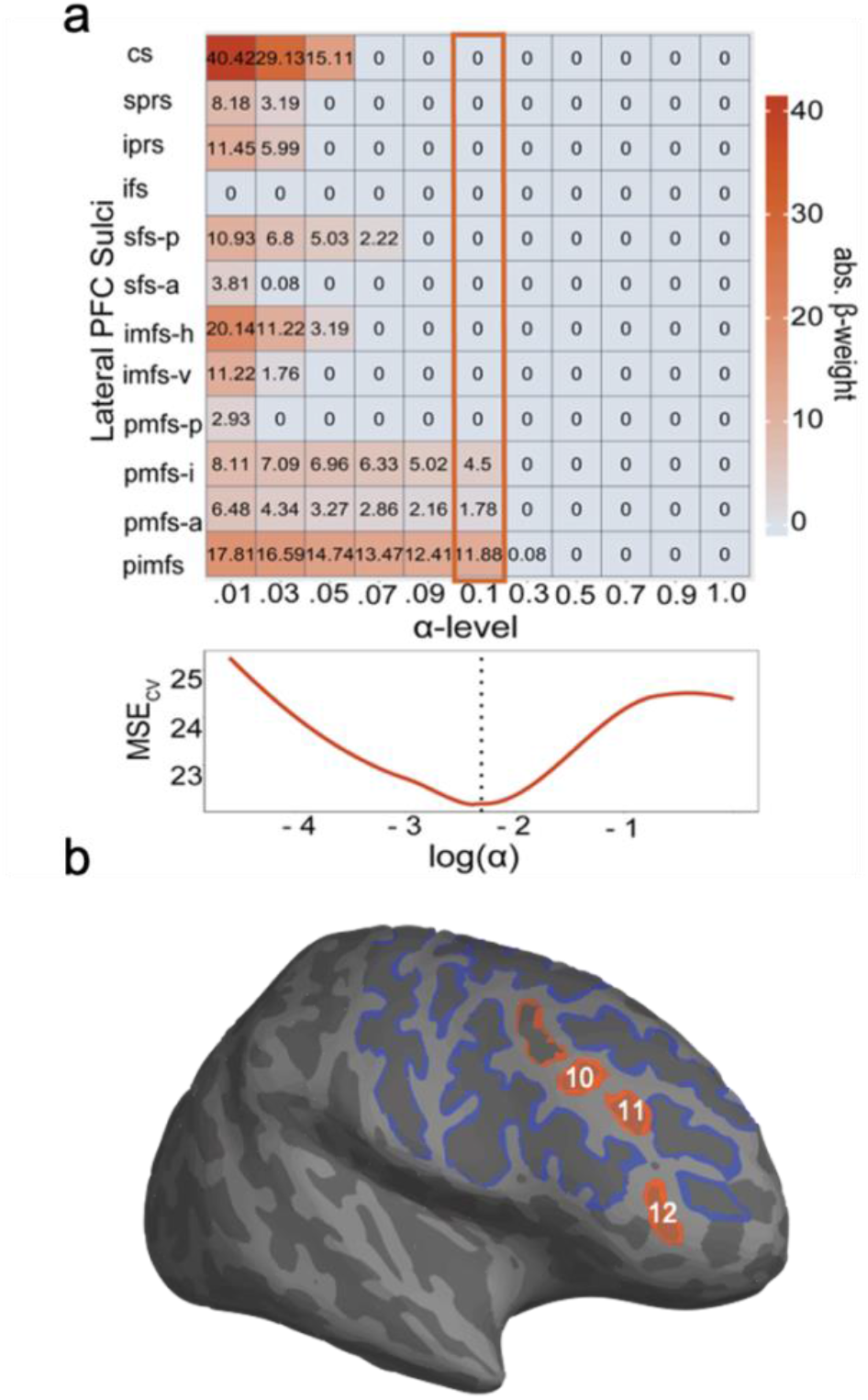
Data-driven model-selection reveals that the depth of a subset of tertiary sulci is associated with reasoning. **a.** Results from the LASSO regression predicting Matrix Reasoning score from sulcal depth in the *Discovery* sample. *Top:* Beta-coefficients for each sulcus at a range of shrinking parameter (alpha) values. Highlighted box indicates coefficients at the chosen alpha-level. *Bottom:* Cross-validated MSE at each alpha-level. By convention^41^, we selected the *α* that minimized the cross-validated Mean Squared Error (MSE_CV_; dotted line). **b**. Inflated cortical surface from an example participant highlighting the three tertiary sulci (*pmfs-i_RH_* (*10*), *pmfs-a_RH_* (*11*), and *pimfs_RH_* (*12*)) implicated in reasoning performance.

To evaluate the generalizability of the sulcal-behavioral relationship identified in the *Discovery* sample, we constructed a linear model to predict reasoning from sulcal depth and age in our *Replication* sample. The mean depths of the *pmfs-i_RH_, pmfs-a_RH_*, and *pimfs_RH_*, as well as age, were included as predictors in the model, as they were the only three sulci identified in the sulcal-behavioral model in the *Discovery* sample. As age was, as expected, highly associated with reasoning (Fig. 5b), including age in this model allowed us to compare performance of this tertiary sulci + age model to a model with age alone in order to determine the unique contribution of LPFC tertiary sulcal depth to reasoning performance above and beyond age. This model (and all subsequent models) were fit using a leave-one-out cross-validation (looCV) procedure. LooCV assesses the generalizability of the model within a sample, however, while appropriate for smaller sample sizes, it can result in models with high variance compared to other cross-validation techniques. To address this concern, we also estimated empirical MSE confidence intervals using a bootstrapping procedure (Online Methods; Fig. 5c). High variance in MSE across the bootstrapped iterations would suggest that the model is likely overfit to the original data.

**Figure 5.**
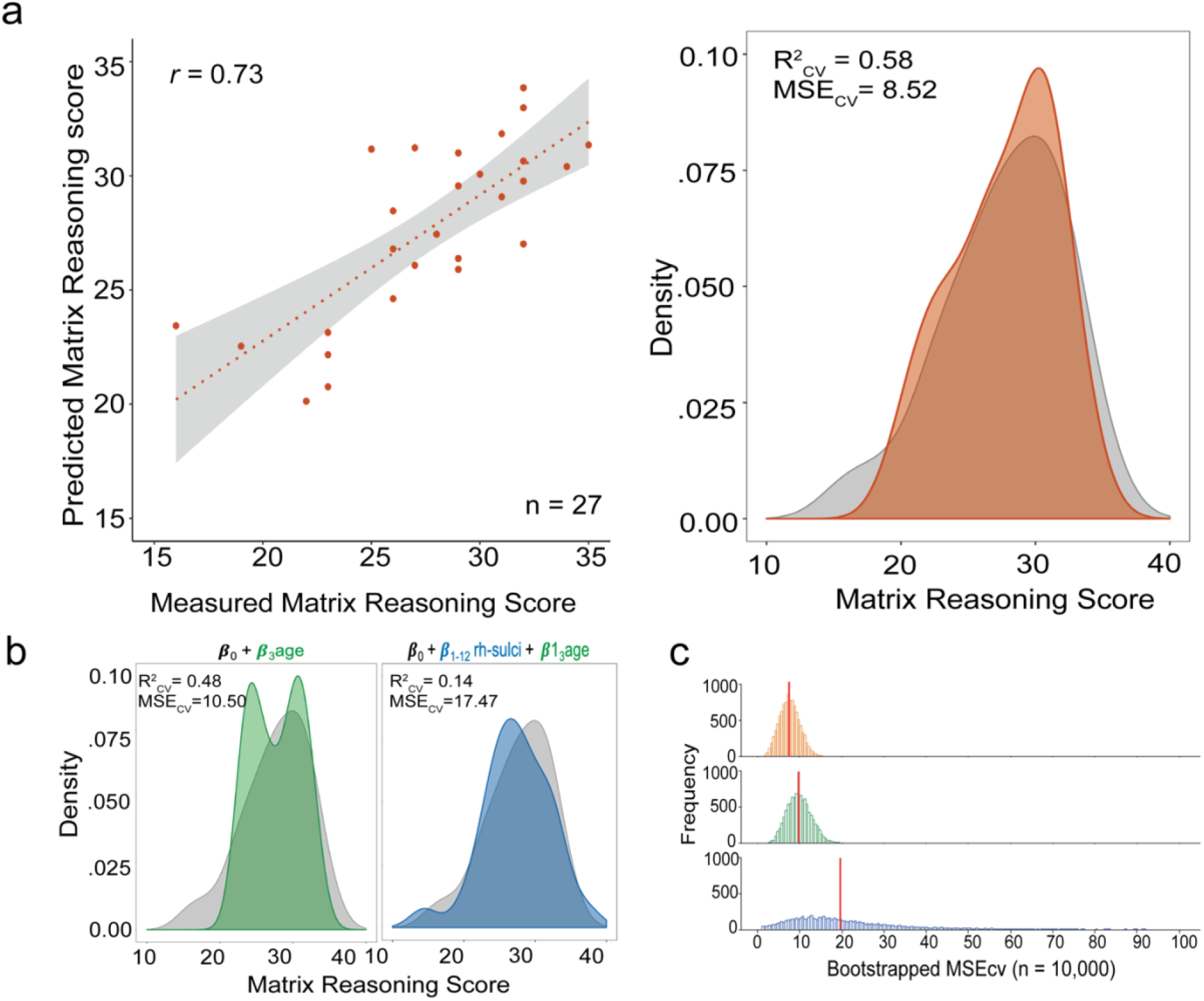
A model-based approach with nested cross-validation reveals that the depth of a subset of LPFC tertiary sulci predicts individual variability in reasoning above and beyond age. **a.** *Left:* Spearman’s correlation between measured and predicted Matrix Reasoning scores in the *Replication* sample for the best tertiary sulci + age model, which includes the depths of the two most predictive sulci (*pmfs-i_RH_* + *pimfs_RH_*) from the *Discovery* sample, as well as age (Supplementary Fig. 5 for a model with all three tertiary sulci selected from the *Discovery* sample). *Right:* Density plot of model fit. The predicted scores from the chosen model (*pmfs-i_RH_* + *pimfs_RH_* + age) are shown in orange and overlaid on the distribution of measured Matrix Reasoning scores (*gray*). **b.** Distribution of predicted scores for the cross-validated nested model comparisons. *Green*: age only. *Blue*: all RH LPFC sulci + age. Each of the model fits are overlaid on the distribution of measured Matrix Reasoning scores (*gray*). The *pmfs-i_RH_* + *pimfs_RH_* + age model (**a**) produced a better fit than both comparison models. **c.** Empirical MSE for each of the three models estimated with a bootstrapping procedure (n_iterations_ = 10,000) to address the potential for looCV to result in high variance and overfitting. The model including all LPFC RH sulci + age (*blue*) exhibited notably high variance in error estimation. The red vertical line indicates the estimated median MSE.

We found that this model (*pmfs-i_RH_* + *pmfs-a_RH_* + *pimfs_RH_* + age) was highly predictive of reasoning score in the *Replication* sample (R^2^_cv_= 0.52, MSE_CV_ = 9.66; Bootstrapped 95%CI_MSE_: 3.12-13.69, median_MSE_ = 8.14). Additionally, we observed a high correspondence (Spearman’s *rho* = 0.70) between predicted and actual measured reasoning scores (Supplementary Fig. 5). Furthermore, if we consider just the two LPFC tertiary sulci that are the strongest predictors of reasoning performance as identified in the *Discovery* sample (*pmfs-i_RH:_ β*_pmfs-i_ = 4.50; *pimfs_RH_: β*_pimfs_ = 11.88), the predictions of reasoning performance and model fits improved even further in the *Replication* sample (R^2^_cv_= 0.58; MSE_CV_ = 8.52; Bootstrapped 95%CI_MSE_ = 3.21-12.37, median_MSE_ = 7.47; Spearman’s *rho* = 0.73; Fig. 4a).

Once we had determined that the sulci relevant for reasoning in the *Discovery* sample were also predictive of reasoning in the *Replication* sample, we used cross-validation to evaluate the fit of the replication model relative to two alternative models considering either 1) age alone or 2) sulcal depth from all right hemisphere LPFC sulci and age together in the *Replication* sample (Fig. 1d). This nested model comparison allowed us to determine the unique contribution of tertiary sulcal depth while still accounting for the effects of age and primary LPFC sulcal depth on reasoning performance. Removing the *pmfs-i_RH_*, *pmfs-a_RH_*, and *pimfs_RH_* from the model decreased prediction accuracy and increased the MSE_cv_ (R^2^_cv_ = 0.48, MSE_CV_ = 10.50; Bootstrapped 95%CI_MSE_ = 4.69-15.67, median_MSE_ = 9.66), indicating that the depths of these right hemisphere tertiary sulci uniquely contribute to the prediction of reasoning scores above and beyond age (Fig. 5b). Additionally, considering age and the depths of all RH LPFC sulci also weakened the model prediction and increased MSE_cv_ (R^2^_cv_ = 0.14, MSE_CV_ = 17.47, Bootstrapped 95%CI_MSE_ = 2.79-306.25, median_MSE_ = 19.70). The bootstrapped CI_MSE_ showed that this model also suffered from very high variance (Fig. 5c). Taken together, our cross-validated, nested model comparison empirically supports that the depth of only a subset of LPFC tertiary sulci reliably explains unique variance in reasoning performance that is not accounted for by age or the depths of all LPFC sulci.

Finally, while our data demonstrate support for our hypothesis, we wondered whether our findings extended to other neuroanatomical features or related measures of cognitive development. We repeated our procedure with 1) a model in which we replaced sulcal depth with cortical thickness^61–64^ and 2) a model in which we replaced reasoning performance with performance on a behavioral measure that reflects a general cognitive ability: processing speed^65^. We used the Akaike Information Criterion (AIC) to quantitatively compare models. If the Δ*AIC* is greater than 2, it suggests an interpretable difference between models. If the Δ*AIC* is greater than 10, it suggests a strong difference between models, with the lower AIC value indicating the preferred model^66,67^ (Online Methods).

With respect to extension of these findings to another anatomical feature, this approach revealed that a model with cortical thickness and age was predictive of reasoning (R^2^_cv_ = 0.33; MSE_CV_ = 13.54), but much less than the model with age alone (R^2^_cv_ = 0.48; MSE_CV_ = 10.50). The AIC for the thickness + age model (AIC_Thickness_ = 78.58) was much higher than the AIC for the tertiary sulci + age model (AIC_sulcalDepth_ = 63.85; ΔAIC_Thickness-Depth_ = 14.73). This indicates that sulcal depth is strongly preferred as a predictor over cortical thickness (Supplementary Fig. 6a),

To test whether sulcal depth would predict another cognitive measure aside from reasoning, we used a test of processing speed (Cross-Out^38^; Fig. 1b). Processing speed is a general cognitive ability that is correlated with—and theorized to support—reasoning^28,31,65,68^. As predicted based on the prior literature, processing speed was correlated with reasoning performance in our sample (*rho* = 0.54, Supplementary Fig. 6c). Sulcal depth of the three critical LPFC tertiary sulci (*pmfs-i_RH_, pmfs-a_RH_*, and *pimfs_RH_*) and age was predictive of processing speed (R^2^_cv_= .45; MSE_CV_=20.53), but not much more than age alone (R^2^ = 0.42; MSE_CV_ = 21.82). The AIC for the processing speed + age model (AIC_cross-Out_ = 89.59) was much higher than the AIC for the tertiary sulci + age model (AIC_sulcalDepth_ = 63.85; ΔAIC_cross-Out-MatrixReasoning_ = 25.74), which indicates that reasoning is strongly preferred over processing speed (Supplementary Fig. 6b).

To further probe the relationship between these sulci and reasoning, we performed a follow-up analysis with a measure of phonological working memory (Forwards Digit Span) as another point of comparison. Like processing speed, working memory is a general cognitive ability that is correlated with—and theorized to support—reasoning^28,69^. As predicted based on the literature, our measures of reasoning and working memory were correlated (*rho* = 0.58; Supplementary Fig. 6c). However, our model did not predict working memory (R^2^_cv_ = 0.10, MSE_cv_ = 2.75).

### Probability maps of LPFC sulci in a pediatric cohort

As this is the first developmental dataset of tertiary sulci in the frontal lobe, we sought to generate spatial probability maps that can be shared with the field. The benefit of such maps is that they capture both the stable and variable features of these sulci across participants. We calculated probability maps^19^ across all participants with at least one identifiable *pimfs* component in each hemisphere (N=58). We provide examples of the un-thresholded probability maps which capture the spatial variability across participants, as well as maps thresholded at 20% and 33% overlap across participants (Fig. 6a). Thresholding captures the shared features across participants and can be applied to increase the interpretability and reduce spatial overlap between sulci^19^ (Online Methods). These probability maps can be projected to cortical surfaces in individual participants across ages (Fig. 6b) and can guide future research that aims to shed light on how LPFC tertiary sulcal morphology affects the functional organization in LPFC, as well as cognition.

**Figure 6.**
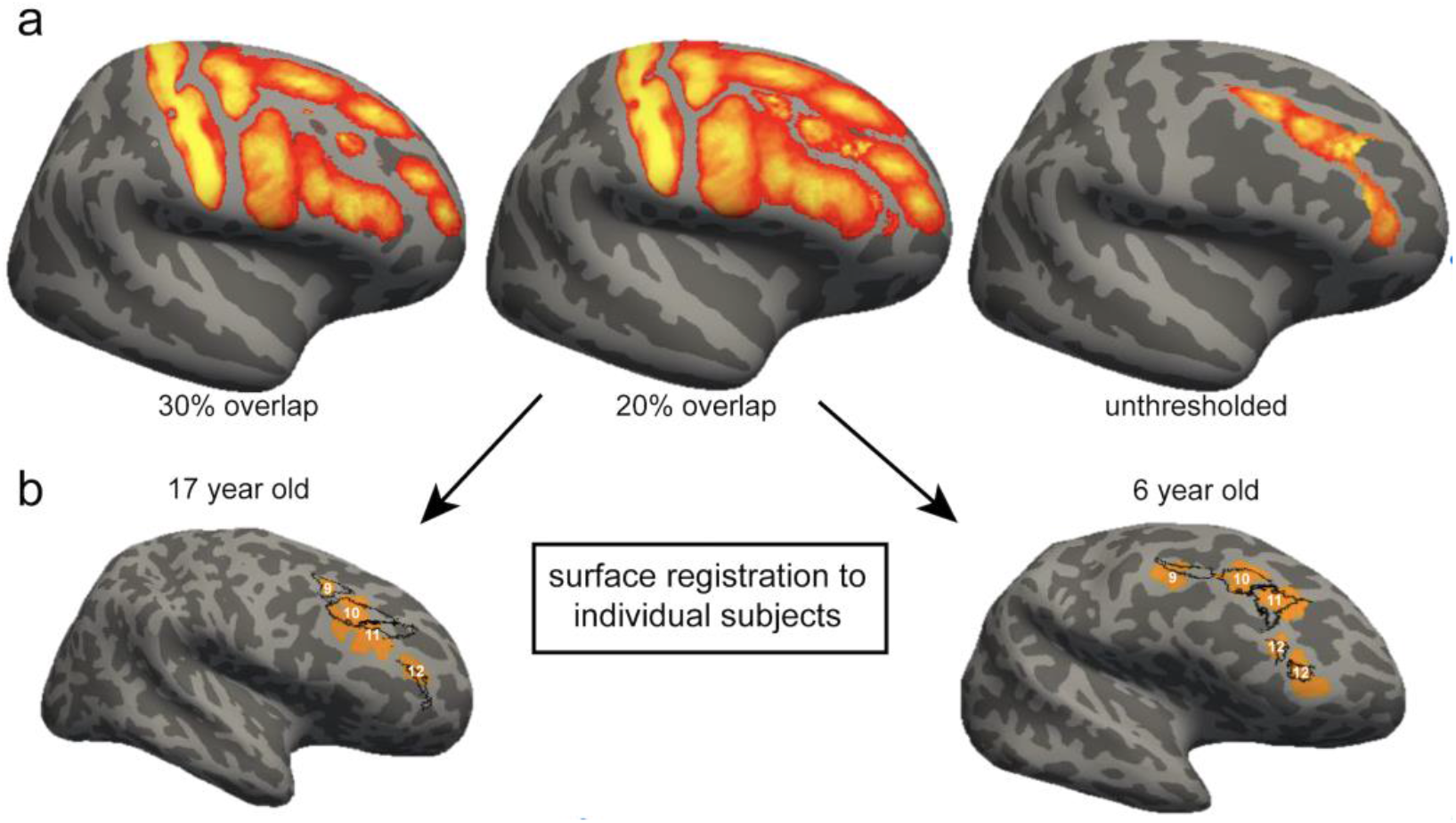
Probability maps of LPFC tertiary sulci. **a.** Maximum probability maps^19^ were generated across all participants with an identifiable *pimfs* in both hemispheres (N=58). To generate the maps, each label was transformed from individual subject space to the common *fsaverage* space. For each vertex, we calculated the proportion of participants for whom that vertex is labeled as the given sulcus. In the case of multiple labels for one vertex, the sulcus with the highest subject overlap was assigned to a given vertex. To reduce spatial overlap, these maps can be thresholded to only include vertices with a minimum percent overlap across participants (eg. 33% (left) or 20% (middle) overlap). The un-thresholded maps (right) can be used to visualize the spatial variability regarding the sulci of interest. **b.** Maps can be projected to individual subject space to guide the definition of tertiary sulci in LPFC. Here, the thresholded maps (20%) for the tertiary sulci are projected back to example (randomly chosen) hemispheres from a 17 year-old (left) and a 6 year-old (right). The outline of the spatial probability maps (black) are overlaid on the manual sulcal definitions (orange) for visualization purposes. While there is not a perfect correspondence between the maps and the tertiary sulci, the maps can guide manual definitions performed by researchers interested in examining LPFC tertiary sulci in future studies. These maps can be applied to other samples and can be downloaded from https://github.com/cnl-berkeley/stable_projects/tree/main/CognitiveInsights_SulcalMorphology. 9: pmfs-p; 10: pmfs-i; 11: pmfs-a; 12: pimfs.

## Discussion

Recent studies examining sulcal morphology in humans and other species continue to improve our understanding of the development and evolution of association cortices. They also provide anatomical insights into cognitive skills that set humans apart from other species^1,19,57,70^. A consistent finding from these previous studies is that developmentally and evolutionarily meaningful changes in sulcal morphology are not homogeneous within association cortices; instead, such changes are focal and related to different aspects of neuroanatomical and functional networks that are behaviorally meaningful^19,39,44,71–76^. After manually defining 1,320 sulci in individual participants and implementing a data-driven approach with nested cross-validation in both *Discovery* and *Replication* samples, our results are consistent with and extend these previous findings by showing for the first time that the sulcal depth of three LPFC tertiary sulci predict behavioral performance on a reasoning task in a developmental cohort above and beyond age. In the sections below, we discuss 1) potential underlying mechanisms that likely contribute to the relationship between tertiary sulcal depth and cognitive performance, 2) how the present findings provide a foundation for future studies attempting to link the morphology of brain structures to behavior and functional brain representations, and 3) how our novel, model-based approach can be applied to study other association cortices across the lifespan.

An immediate question generated from our findings is: *What underlying mechanisms could explain why the depths of LPFC tertiary sulci and age reliably predict reasoning performance on a complex behavioral task?* We offer one potential explanation that integrates recent anatomical findings^19,77^ with a classic theory^9^ and propose a hypothesis linking sulcal depth to short-range anatomical connections, and in turn, to cortical networks and cognitive performance. Specifically, in the 1960s, Sanides^9,22^ proposed that morphological changes in tertiary sulci would likely be associated with the development of higher order processing and cognitive skills. The logic of Sanides’ hypothesis extends from the fact that tertiary sulci emerge last in gestation and have a protracted development after birth, while complex cognitive skills such as reasoning ability also have a protracted development in childhood. Our findings support this classic hypothesis. However, while the LPFC is considered critical to reasoning^6,24,30,32,33^, reasoning performance cannot be localized to a single structure^6,24,78,79^ and thus, the mechanism behind this relationship still needs to be investigated.

As a starting point toward understanding the underlying mechanism, two recent empirical findings provide underlying anatomical mechanisms that could support this relationship between tertiary sulci and cognition. First, there is a relationship between human LPFC tertiary sulcal morphology and myelination^19^, which is critical for short- and long-range connectivity, as well as the efficiency of communicating neural signals among regions within cortical networks^80^. Second, anatomical work in non-human primates has shown that long-range white matter fiber tracts have a bias for terminating in gyri, while additional short-range white matter fibers commonly project from the deepest points (*fundi*) of sulci^77^, which we refer to as *fundal fibers*. These previous and present findings serve as the foundation for the following novel mechanistic hypothesis linking tertiary sulcal depth to anatomical connections and neural efficiency: deeper tertiary sulci likely reflect shorter fundal fibers, which in turn, reduce the length of short-range anatomical connections between cortical regions, and thus, increase neural efficiency. While speculative, this hypothesis is similar in logic to the tension-based theory of cortical folding^81^ and also feasible given the fact that short-range structural connectivity increases and sulci deepen during development^82,83^. This increase in neural efficiency could underlie variability in cognitive performance, which can be tested in future studies incorporating anatomical, functional, and behavioral measures, as well as computational modeling.

In addition to this mechanistic hypothesis, our present findings improve the spatial scale of previous studies attempting to link cortical morphology to behavior associated with LPFC. For example, previous studies identified an association between cognitive skills and cortical thickness of LPFC in its entirety^61–64^. While we find an association between reasoning and cortical thickness, when considering individual tertiary sulci, our analyses indicate that the depths of tertiary sulci and age together are much stronger predictors of reasoning than the cortical thickness of these sulci and age together. In fact, when including the cortical thickness of sulci in the model, performance is worse than age alone (Supplementary Fig. 6a). The combination of these findings across studies suggests that neuroanatomical-behavioral relationships can exist at multiple spatial scales in the same macro-anatomical expanse such as LPFC: cortical thickness at the macroanatomical scale and tertiary sulcal depth at the meso-scale.

We also emphasize that, though our model-driven approach identified a subset of LPFC tertiary sulci, morphological features of these sulci are likely correlated with other cognitive tasks, and it is highly probable that other LPFC tertiary sulci play critical roles in other tasks beyond reasoning. Although we did not observe a relationship between the depth of these sulci and two related cognitive measures, this should not be taken as evidence that these sulci show specificity to reasoning. We also emphasize that the present approach of precise anatomical mapping of tertiary sulci does not imply that reasoning can be localized to a single sulcus, or even a single cortical region. In fact, our previous work, including previous studies on this dataset, has focused extensively on the distributed nature of reasoning, highlighting patterns of functional and structural connectivity between prefrontal and parietal regions that support this process^6,24,30^. Additionally, focusing on tertiary sulci in PFC forms a foundation for understanding how these largely overlooked neuroanatomical structures contribute to typical brain function and cognition, especially at the network level^45^. Indeed, modern multi-modal neuroimaging research from two recent parallel lines of work show that meticulously labeling tertiary sulci within individuals uncovers new structural-functional relationships within PFC at the network level^19,57^. Thus, future studies exploring the relationship between sulcal morphology and behavioral performance in additional cognitive tasks at the level of individual participants will begin to generate a more comprehensive sulcal-behavioral map in LPFC with additional insights into cortical networks.

In addition to this sulcal-behavioral map in LPFC, two recent lines of work show feasibility for future studies attempting to link tertiary sulcal morphology to brain function, especially for functional activity related to reasoning: one related to tertiary sulci as a meso-scale link between microstructural and functional properties of LPFC and the other identifying functional representations related to reasoning. In terms of the former, a series of recent studies have shown that tertiary sulci are critical functional landmarks in different association cortices^13,39,58,12^ and variability in sulcal morphology in the medial prefrontal cortex has been associated with changes in cortical morphometry linked to individual differences in cognitive performance and clinical symptom presentation in schizophrenia^46^. Additionally, in LPFC, Miller and colleagues^13^ showed that the different *pmfs* components explored here were functionally distinct in adults with respect to resting-state connectivity profiles. In terms of the latter, numerous functional neuroimaging studies show that LPFC is central for reasoning performance^32,84^. More explicitly, several studies also indicate that the middle frontal gyrus, the gyrus in which the three sulci (*pmfs-i, pmfs-a*, and *pimfs*) identified by our model are located, plays an important role in cognitive processes that are integral for reasoning, such as maintaining representations and forming associations^3,4^. Thus, future investigations of functional connectivity, as well as functional representations, relative to tertiary sulci in future studies in children and adults will likely bring us closer to understanding the complex relationship between the development of LPFC anatomical organization, functional organization, and behavior.

While we limit our focus to the LPFC in the present study, both because of its relevance for reasoning, but also because of the immense manual labor involved in this type of study, the novel, data-driven pipeline introduced here can be applied to any cortical expanse. For example, lateral parietal cortex is also critical for relational reasoning, is expanded in humans compared to non-human primates^2,85^, and also contains tertiary sulci^20^. Additionally, structural connectivity between frontal and parietal regions increases across development^24,86,87^. Thus, future studies can explore how morphological features of tertiary sulci in a) LPFC and lateral parietal cortex contribute to reasoning performance and b) different association cortices contribute to performance on cognitive tasks, as well as functional representations in each cortical expanse. It will also be important to explore the relationship among tertiary sulci across cortical regions. For example, developmental studies are well suited to explore how the variability in sulcal morphology in one cortical region, such as LPFC, might affect morphology of tertiary sulci in other cortical regions, such as medial frontal or parietal regions. Our modeling approach can also be applied to data across the lifespan – either cross-sectionally or longitudinally. While it is known that tertiary sulci are shallow indentations in cortex that emerge last in gestation (relative to primary and secondary sulci), and have a protracted development after birth^9,11,14,15,20,37,38,57,88^, the history of LPFC sulcal definitions, especially within the MFG, has been contentious^7,8,17–22,9–16,45^. Thus, while we used these classic studies to label the type of each sulcus, the distinctions among primary, secondary, and tertiary sulci should be confirmed by modern studies of cortical folding in gestation. Crucially, our findings are not dependent on this classification. Our data-driven, model-based approach identified that a subset of shallow sulci in LPFC explain the most variance in reasoning skills across participants above and beyond age in both *Discovery* and *Replication* samples. Additionally, the developmental timeline of tertiary sulci relative to the development of functional representations and cognitive skills is unknown. Future studies implementing and improving our model-based approach can begin to fill in these gaps in the developmental timeline of tertiary sulci anatomically, behaviorally, and functionally.

Despite the many positive applications of our model-based approach and the many future studies that will likely build on the foundation of the present novel findings, there are also limitations. The main drawback of the precise, single-subject approach implemented here is that it relies on manual sulcal definitions, which are time-consuming and require anatomical expertise. This limits sample sizes and the expanse of cortex that can be feasibly explored in a given study. Additionally, while there is “no one-size-fits-all sample size for neuroimaging studies”^84^ and we had a large N (>1000) in terms of sulci explored in the present study, new methods and tools will need to be developed to increase the number of participants in futures studies. Increasing the number of participants will improve the diversity of our sample and reduce imbalances in gender or other demographic features. Ongoing work is already underway to develop deep learning algorithms to accurately define tertiary sulci automatically in individual participants, and initial results are promising.^89,90^ In the interim, our probabilistic sulcal maps can guide manual definitions performed by researchers interested in examining LPFC tertiary sulci in future studies (Fig. 6).

In summary, using a data-driven, model-based approach, we provide cognitive insights from evolutionarily new brain structures in human LPFC for the first time. After manually defining 1,320 LPFC sulci, our approach revealed that the depths of tertiary sulci reliably predicted reasoning skills above and beyond age. Methodologically, our study opens the door for future studies examining these evolutionarily new tertiary sulci in other association cortices, as well as improves the spatial scale of understanding for future studies interested in linking cortical morphology to behavior. Theoretically, the present results support an untested anatomical theory proposed over 55 years ago^9^. Mechanistically, we outline a novel hypothesis linking tertiary sulcal depth to short-range white matter fibers, neural efficiency, and cognitive performance. Together, the methodological, theoretical, and mechanistic insights regarding whether, or how, tertiary sulci contribute to the development of higher-level cognition in the present study serve as a foundation for future studies examining the relationship between the development of cognitive skills and the morphology of tertiary sulci in association cortices more broadly.

## Online Methods

### Participants

The present study consisted of *Discovery* (N=33; 16 males and 17 females) and *Replication* (N=28; 20 males and 8 females) samples. For the *Discovery* sample, 33 typically developing individuals between the ages of 6-18 were randomly selected from the Neurodevelopment of Reasoning Ability (NORA) dataset^24,30,31,36^. Following the definition of sulci in this sample, we selected an additional 28 age-matched participants for the *Replication* sample. No features other than age were considered in the selection of the *Replication* sample. The terms *male* and *female* are used to denote parent reported gender identity. All participants were screened for neurological impairments, psychiatric illness, history of learning disability, and developmental delay. All participants and their parents gave their informed assent and/or consent to participate in the study, which was approved by the Committee for the Protection of Human Participants at the University of California, Berkeley.

## Data Acquisition

### Imaging data

Brain imaging data were collected on a Siemens 3T Trio system at the University of California Berkeley Brain Imaging Center. High-resolution T1-weighted MPRAGE anatomical scans (TR=2300ms, TE=2.98ms, 1×1×1mm voxels) were acquired for cortical morphometric analyses.

### Behavioral data

Behavioral metrics are only reported for the participants included in the morphology-behavior analyses (*Discovery*: n = 28, *Replication*: n = 27). Reasoning performance was measured as a total raw score from the WISC-IV Matrix Reasoning task^39^ (Fig. 1b; *Discovery:* mean(sd) = 24.28 (4.86); *Replication:* mean(sd) = 27.64 (4.52)). Matrix Reasoning is an untimed subtest of the WISC-IV in which participants are shown colored matrices with one missing quadrant. The participant is asked to “complete” the matrix by selecting the appropriate quadrant from an array of options (Fig. 1b). Matrix Reasoning score was selected as it is a widely used measure of non-verbal reasoning^30,31^ and it was the most consistently available reasoning measure for the participants in this study. Matrix Reasoning has previously been examined in relation to white matter and functional connectivity in a large dataset that included these participants^30^ and a previous factor analysis in this dataset showed that the Matrix Reasoning score loaded strongly onto a reasoning factor that included three other standard reasoning assessments^31^.

Processing speed was computed from raw scores on the Cross-Out task from the Woodcock-Johnson Psychoeducational Battery-Revised^73^ (WJ-R; Fig. 1b). In this task, the participant is presented with a geometric figure on the left followed by 19 similar figures. The participant places a line through each figure that is identical to the figure on the left of the row (Fig. 1b). Performance is indexed by the number of rows (out of 30 total rows) completed in 3 minutes (*Replication:* Mean(sd) = 22.19 (6.26)). Cross-Out scores are frequently used to estimate processing speed in developmental populations.^91,92^

As an additional measure, working memory (WM) was assessed from raw Digit Span Forward scores (*Replication:* Mean(sd) = 9.03(1.77)). Digit Span Forward scores measure WM maintenance and attention. For each forward trial, participants were presented with a string of numbers by the experimenter and were asked to immediately repeat the numbers in the same order. The task consisted of eight questions with two trials per level (16 total trials). Each question (set of two trials) consisted of a longer string of numbers than the question before. Both Processing speed and Working memory were selected as they are considered related, but separable, measures from Reasoning. We report the Spearman correlation coefficient (*rho*) among each of the three behavioral measures (Supplementary Fig. 6).

## Morphological Analyses

### Cortical surface reconstruction

All T1-weighted images were visually inspected for scanner artifacts. FreeSurfer’s automated segmentation tools^93,94^ (FreeSurfer 6.0.0) were used to generate cortical surface reconstructions. Each anatomical T1-weighted image was segmented to separate gray from white matter, and the resulting boundary was used to reconstruct the cortical surface for each subject^93,95^. Each reconstruction was visually inspected for segmentation errors, and these were manually corrected.

### Manual labeling of LPFC sulci

Sulci were manually defined separately in the *Discovery* and *Replication* samples according to the most recent atlas proposed by Petrides (2019)^20^. This atlas offers a comprehensive schematization of sulcal patterns in the cortex. The LPFC definitions have recently been validated in adults^19^, but to our knowledge, these sulci have never been defined in a developmental sample. 12 LPFC sulci were manually defined within each individual hemisphere in tksurfer^19^ (Fig. 3; Supplementary Fig. 1 for all manually defined sulci in 122 hemispheres). Sulcal depth values are a feature of FreeSurfer’s scale, which can be explored further on their website (https://surfer.nmr.mgh.harvard.edu). Briefly, depth values are calculated based on how far removed a vertex is from what is referred to as a “mid-surface,” which is determined computationally so that the mean of the displacements around this “mid-surface” is zero. Thus, generally, gyri have negative values, while sulci have positive values. Given the shallowness and variability in the depth of LPFC tertiary sulci, some mean depth values extend below zero. We emphasize that this just reflects the metric implemented in FreeSurfer. For example, max depth values are above zero for all sulci (Supplemental Fig. 5b). Manual lines were drawn on the inflated cortical surface to define sulci based on the proposal by Petrides^96^, as well as guided by the *pial* and *smoothwm* surfaces of each individual^19^. In some cases, the precise start or end point of a sulcus can be difficult to determine on one surface^90^. Thus, using the *inflated, pial*, and *smoothwm* surfaces of each individual to inform our labeling allowed us to form a consensus across surfaces and clearly determine each sulcal boundary. Our cortical expanse of interest was bounded by the following sulci: (1) the *anterior* and *posterior* components of the *superior frontal sulcus* (sfs) served as the superior boundary, (2) the *inferior frontal sulcus* (ifs) served as the inferior boundary, (3) the *central sulcus* served as the posterior boundary, and (4) the *vertical and horizontal* components of the *intermediate fronto-marginal sulcus* (imfs) served as the anterior boundary. We also considered the following tertiary sulci: *anterior* (pmfs-a), *intermediate* (pmfs-i), and *posterior* (pmfs-p) components of the *posterior middle frontal sulcus* (pmfs), and the *para-intermediate frontal sulcus* (pimfs)^19,20^. Please refer to Fig. 2a for the location of each of these sulci on example hemispheres and Supplementary Fig. 1 for the location of all 1,320 sulci in all 122 hemispheres. For each hemisphere, the location of each sulcus was confirmed by two trained independent raters (W.V. and J.Y.) and finalized by a neuroanatomist (K.S.W). The surface vertices for each sulcus were then manually selected using tools in *tksurfer* and saved as surface labels for vertex-level analyses of morphological statistics. All anatomical labels for a given hemisphere were fully defined before any morphological or behavioral analyses were performed.

While we could not identify the dorsal and ventral components of the *pimfs* in every hemisphere (Results; Supplementary Table 1), we could identify at least one component of the *pimfs* in each hemisphere in nearly all participants in the *Discovery* (28/33) and *Replication* (27/28) samples. Thus, our inclusion criteria for all subsequent analyses was to include participants who had at least one *pimfs* component in each hemisphere, which assures that all repeated measures statistics are balanced for effects of sulcus and hemisphere. For those participants who had identifiable dorsal and ventral *pimfs* components, we merged the components into one label, using the FreeSurfer function *mris_mergelabels* and all findings are reported for the merged label^93^.

### Characterization of tertiary sulcal patterning

For each tertiary sulcus, we characterized sulcal patterns, or types, based on intersections with surrounding sulci. We report the number of intersections for a given sulcus with every other sulcal pair (except the central sulcus as no tertiary sulcus intersected with the central sulcus), relative to the total frequency of occurrence of that sulcus in the hemisphere (Figure 2b). We report correlations between left and right hemispheres in each sample as well as the correlation between samples.

### Sulcal probability maps

Sulcal probability maps were calculated to describe the vertices with the highest and lowest correspondence across participants^19^. These maps were generated across all participants with at least one *pimfs* component in each hemisphere. To generate the maps, each label was transformed from individual subject space to the common *fsaverage* space. We chose to use the standard *fsaverage* template to increase accessibility for future studies. Then, for each vertex, we calculated the proportion of participants for whom that vertex is labeled as the given sulcus. in the case of multiple labels, labels were assigned to each vertex with a “winner-take-all” approach. That is, the sulcus with the highest subject overlap was assigned to a given vertex. Consistent with Miller et al. (2021)^19^, in addition to providing un-thresholded maps, we also constrained these maps to be maximum probability maps (MPMs) which helps avoid overlapping sulci and can increase interpretability (Fig. 6a)^19^. We provide thresholded maps at 33% and 20% spatial overlap for each label. This allows the user to assess both the spatial variability between participants as well as the stable features shared across participants. Finally, since this is the first developmental dataset of tertiary sulci in the frontal lobe, we make these maps publicly available for download: https://github.com/cnl-berkeley/stable_projects/tree/main/CognitiveInsights_SulcalMorphology.

### Characterization of sulcal morphology

As the most salient morphological feature of tertiary sulci is their shallowness compared to primary and secondary sulci^9,11,14,15,20,23,37,39^, we focused morphological analyses on measures of sulcal depth. Raw depth metrics (standard FreeSurfer units) were computed in native space from the .sulc file generated in FreeSurfer 6.0.0^93^. We normalized sulcal depth to the maximum depth value within each individual hemisphere in order to account for differences in brain size across individuals and hemispheres. All depth analyses were conducted for normalized mean sulcal depth. As cortical thickness is a commonly used metric in developmental studies, we also considered the mean cortical thickness (mm) for each sulcus. Mean cortical thickness for each sulcal label was extracted using the *mris_anatomical_stats* function that is included in FreeSurfer^94^.

### Distinction among primary, secondary, and tertiary sulci

As described in our previous work, as well as classic studies, tertiary sulci are defined as the last sulci to emerge in gestation after the larger and deeper primary and secondary sulci^7,8,17–21,9–16,45^ (Fig. 2a). Specifically, previous studies specify that a) primary sulci emerge prior to 32 weeks in gestation, b) secondary sulci emerge between 32-36 weeks in gestation, and c) tertiary sulci emerge during and after 36 weeks.^97^ Previous research identifies the *cs, prs, sfs, and ifs* as primary sulci. As such, we apply these definitions to the subcomponents of the sfs (*sfs-a and sfs-p*) and prs (*sprs and iprs*) considered here. The *imfs-v* and *imfs-h* are contemporary labels for classic definitions of sulci commonly labeled as either the *frontomarginal* and/or *middle frontal sulci*^19,20,45^. When considering classic papers and atlases^10,16,18,21^ both the *imfs-h* and *imfs-v* appear to be prevalent prior to 32 weeks. Thus, we classify them as primary sulci in the present study (see also Miller et al., 2021b)^45^. Finally, while classic analyses have not considered modern definitions of *pmfs* and *pimfs* sulcal components, we refer to them as tertiary sulci for two main reasons. First, from our historical analyses, sulci within the middle frontal gyrus emerge during late stages in gestation, consistent with definitions of tertiary sulci^45^. Second, from our previous analyses in adults^19^, *pmfs* sulcal components are small and shallow relative to other primary and secondary LPFC sulci, which is consistent with morphological features of tertiary sulci. Taken together, our distinction among primary and tertiary sulci is based on classic and modern data. Future studies with larger sample sizes using non-invasive fetal imaging will re-visit the timestamps for the documented sulci, as well as provide new timestamps for those sulci that were not included in these classic studies. For example, based on these classic definitions of sulcal types, the present study did not include any secondary sulci in LPFC. Nevertheless, we also highlight that our data-driven approach is blind to these definitions and identifies sulci that are small in surface area and shallow in depth, which is consistent with the definition of tertiary sulci.

### Comparison between tertiary and primary sulci

We compared sulcal depth of tertiary and primary sulci with a 2-way (hemisphere x sulcal type) repeated measures analysis of variance (rm-ANOVA; Fig. 3c). To assess the variability in depth between hemispheres and groups, we conducted the same rm-ANOVA, but replaced mean sulcal depth with the standard deviation. We conducted the same repeated measures analyses with cortical thickness between tertiary and primary sulci in both samples (Supplementary Fig. 3; see Supplementary results). All ANOVAs were computed in R with the *aov* function, imported in python via *rpy2*. Effect sizes are reported with the *generalized* eta-squared (*η*^2^) metric.

## Assessing the relationship between sulcal depth and reasoning performance

### Four-pronged analytic approach

Based on current recommendations^59^, we implement a four-pronged approach to assess and improve the generalizability of our findings at each stage of analysis.

1. <u>Regularization:</u> In the Discovery sample, we use L1-regularization (LASSO-regression) as part of our model selection approach. Not only does this provide a data-driven method for model selection, but regularization techniques are recommended to improve the generalizability of a model^59,60^. Unlike many techniques that only *assess* generalizability, L1 regularization actually *increases* the generalizability of a model by providing a sparse solution that reduces coefficient values and decreases variance in the model without increasing bias. This technique guards against overfitting and increases the likelihood that a model will generalize to other datasets.
2. <u>Cross-validation:</u> In addition to using regularization techniques to improve generalizability, all our models are fit with cross-validation. The purpose of crossvalidation is to *test* the generalizability of a model within a sample^53^. We report a very strong model fit for our cross-validated models.
3. <u>Replication in an additional sample:</u> We *demonstrate* the generalizability of our findings by showing that the depths of sulci that are predictive of reasoning in the *Discovery* sample generalize to the *Replication* sample. Our regularized regression reveals that the depths of a subset of RH tertiary sulci are relevant for reasoning performance in the *Discovery sample* (Fig. 3). We then show that these same sulci can be used to predict reasoning with high accuracy in the *Replication sample* (Fig 4a).
4. <u>Bootstrapped error estimates:</u> We use bootstrapping as a diagnostic tool to assess the generalizability of our models to out-of-sample data. Using 10,000 iterations, we show our chosen models have low variance in estimated error (Fig. 4d), suggesting that they are not over-fit to the data, and the findings will likely generalize to other samples.

### Model selection - Discovery sample

We applied a least absolute shrinkage and selectionoperator (LASSO) regression model to determine which sulci, if any, were associated with Matrix Reasoning^39^. The depth of all 12 LPFC sulci were included as predictors in the regression model. This analysis was performed separately for each hemisphere. LASSO performs L1-regularization by applying a penalty, or shrinking parameter(*α*), to the absolute magnitude of the coefficients such that:

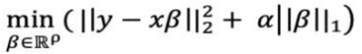

In a LASSO regression, low coefficients are set to zero and eliminated from the model. In this way, LASSO can facilitate variable selection, leading to simplified models with increased interpretability and prediction accuracy^60^. In our case, the LASSO regression algorithm shrinks the coefficients of each of the sulci until only the sulci most predictive of reasoning remain in the model. The LASSO regression model was conducted separately for left and right hemispheres. By convention, we used cross-validation to select the shrinking parameter (*α*). We used the SciKit-learn GridSearchCV package^98^, to perform an exhaustive search across a range of *α*-values (0.01-10.0), and selected the value that minimized cross-validated Mean Squared Error (MSE_CV_).

### Model evaluation - Replication sample

To further characterize the relationship between sulcal depth and reasoning performance, we used the predictors identified by the LASSO-regression in the *Discovery* sample to predict Matrix Reasoning score in the *Replication* Sample. As age is correlated with Matrix Reasoning score, we included age as an additional covariate in the model [1a]. We fit this model as well as alternate nested models with leave-one-out cross validation (looCV). We used nested model comparison to assess the unique variance explained by sulcal depth, while accounting for age-related effects on reasoning:

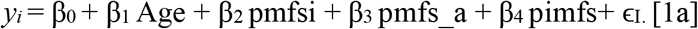

Additionally, we conducted this analysis with only the two most predictive sulci (*pmfs-i, pimfs*) from the *Discovery* sample:

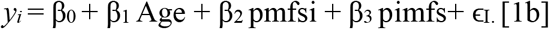

To assess the unique variance explained by tertiary sulcal depth, we compared the MSE_CV_ of this model to the MSE_CV_ of a model with age as the sole predictor [2]:

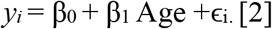

As these models are nested (all predictors in the smaller model [2] are also included in the larger models [1a-b]), we are able to directly compare the prediction error in these two models. Finally, to assess the specificity of the relationship to tertiary sulci in our *Replication* sample, we assessed the fit of model [1] to a full model that included all identified LPFC sulci within a hemisphere [3]. The full model is as follows:

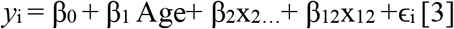

where *x*_2_…*x*_12_ represent the sulcal depth of each identified sulcus within a hemisphere.

### Empirical MSE confidence intervals

The size (n = 27) of the *Replication* sample makes looCV suitable. However, models that are fit with looCV can have high variance. Thus, to assess the potential variance in our estimations, we performed a bootstrapping procedure to empirically estimate the distribution of possible MSE_cv_ predictions for models 1b, 2, and 3. For each model, data were randomly selected with replacement 10,000 times and MSE_cv_ was computed for each iteration. From this process, we estimate Median MSE and 95% confidence intervals for each model (shown in Fig. 3d). All analyses were conducted with SciKit-Learn package in Python^77^.

## Assessing morphological and behavioral preference of the model

### Cortical thickness

To assess whether our findings generalized to other anatomical features, we considered cortical thickness, which is an anatomical feature commonly explored in developmental cognitive neuroscience studies^64,99–101^. To do so, we replaced sulcal depth with cortical thickness as the predictive metric in our best performing model in the *Replication* sample [Model 1b]. As with depth, the model was fit to the data with looCV. To compare the thickness model to the depth model, we used the Akaike Information Critertion (AIC), which provides an estimate of in-sample prediction error and is suitable for non-nested model comparison. AIC is given by:

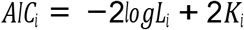

Where *L_i_* is the likelihood for the model (*i*) and *K_i_* is the number of parameters. By comparing AIC scores, we are able to assess the relative performance of the two models. If the Δ*AIC* is greater than 2, it suggests an interpretable difference between models. If the Δ*AIC* is greater than 10, it suggests a strong difference between models, with the lower AIC value indicating the preferred model^66,67^.

### Processing Speed and Working Memory

To ascertain whether the relationship between sulcal depth and cognition is specific to reasoning performance, or transferable to other general measures of cognitive processing^92^, we investigated the generalizability of the sulcal-behavior relationship to two other widely used measures of cognitive functioning: Processing speed and working memory. Specifically, we used looCV to predict processing speed (as indexed by Cross-Out score) and working memory (as indexed by Forwards Digit Span score)^102^ instead of Matrix Reasoning score. In the cases in which the model was predictive, we used AIC to compare the predictions to Matrix Reasoning predictions.

## Supplementary Information

### No differences in cortical thickness among primary and tertiary sulci in LPFC

As discussed throughout this paper, sulcal depth is the main morphological feature differentiating tertiary from primary and secondary sulci^11,14,15,19,20,23,103,104^. Nevertheless, previous developmental work on structural variability in PFC has frequently focused on cortical thickness^64,99–101^. Thus, we also investigated variability in cortical thickness in both the *Discovery* and *Replication* samples as a function of sulcal type (*tertiary* vs. *primary*) and hemisphere (*left* vs. *right*). There were no significant differences in cortical thickness between primary and tertiary sulci in the *Discovery* (F(1,27) = 2.44, *p* = 0.13; Mean(sd))_Tertiary_ = 2.41(0.36); Mean(sd)_Primary_ =2.37 (0.26); Supplementary Fig. 3a) or *Replication* (F(1,26) = 2.31, p =0.14; Mean(sd))_Tertiary_ = 2.31(0.41), Mean(sd)_Primary_ = 2.38(0.30); Supplementary Fig. 3b) samples. Interestingly, the rm-ANOVA revealed a main effect of hemisphere in both samples in which right hemisphere sulci were cortically thinner than left hemisphere sulci (*Discovery:* (F(1,27) = 123.1, *p*<10^-3^, *η*^2^_G_ = 0.09; Mean(sd)_RH_; = 2.30 (0.28); Mean(sd)_LH_ = 2.47(0.27); *Replication:* (F(1, 26) = 42.91, *p*<10^-3^, *η*^2^_G_=0.06; Mean(sd)_RH_ = 2.20(0.36), Mean(sd)_LH_ =2.41(0.29); Supplementary Fig. 3b). Thus, while previous developmental work on structural variability in PFC has focused on cortical thickness^64,99–101^, when considering tertiary sulci, the present analyses emphasize the utility of sulcal depth, not cortical thickness, for differentiating tertiary from primary sulci.

**Supplementary Fig. 1.**
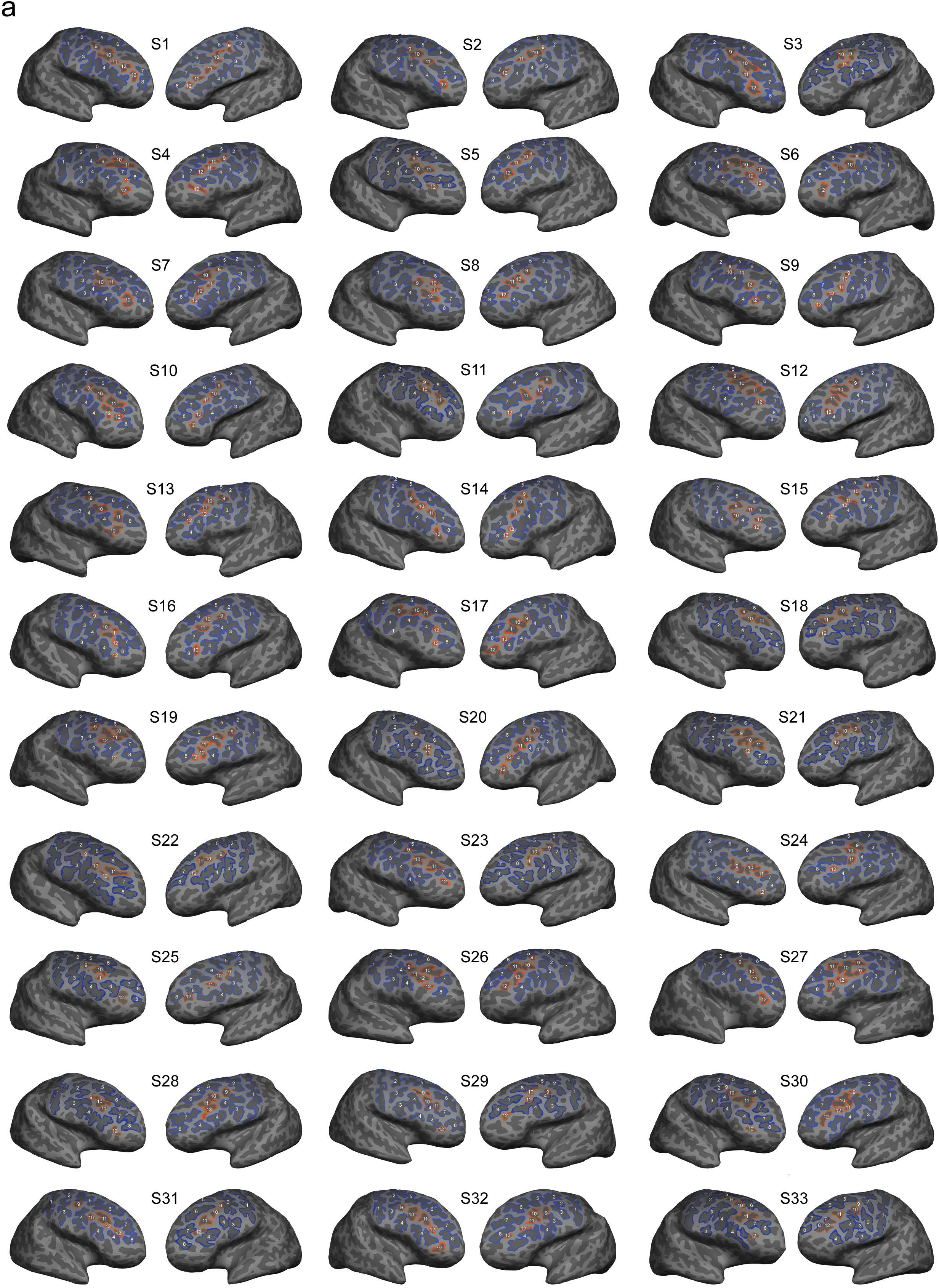
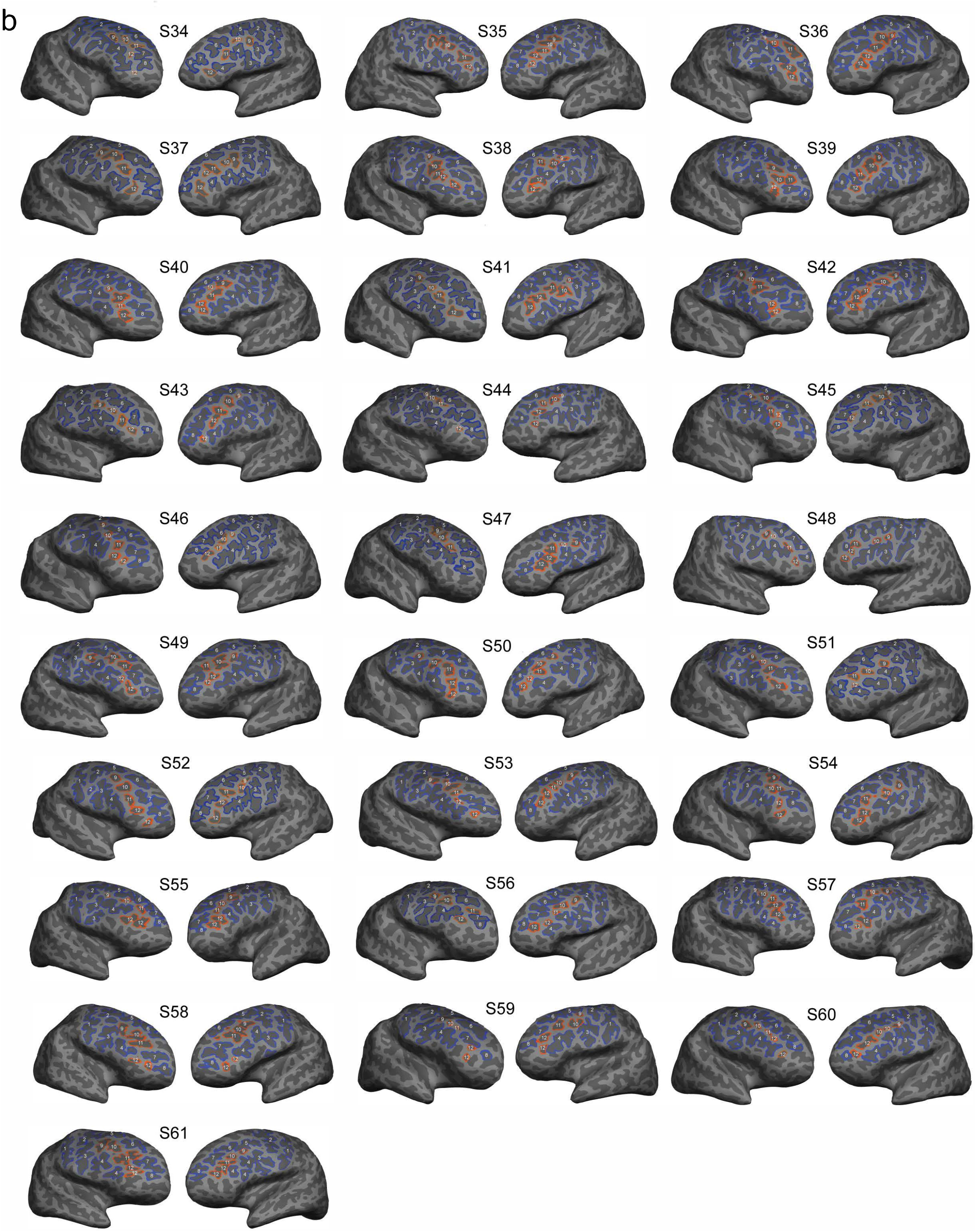
Manual labels in the left and right hemispheres of every subject displayed on the inflated cortical surface. **a.** Manually labeled sulci on the inflated cortical surface in the left and right hemisphere for every subject in the Discovery Sample. Tertiary (orange) and primary/secondary (blue) sulci are identifiable in every subject. The pimfs (*12*) can contain two components, one component, or be absent altogether in a given hemisphere (Supplementary Table 1). **b.** Same layout as in a., but for the Replication Sample. As in the Discovery Sample, tertiary (orange) and primary/secondary (blue) sulci are identifiable in every subject in which the pimfs could contain 0, 1, or 2 components.

**Supplementary Fig. 2.**
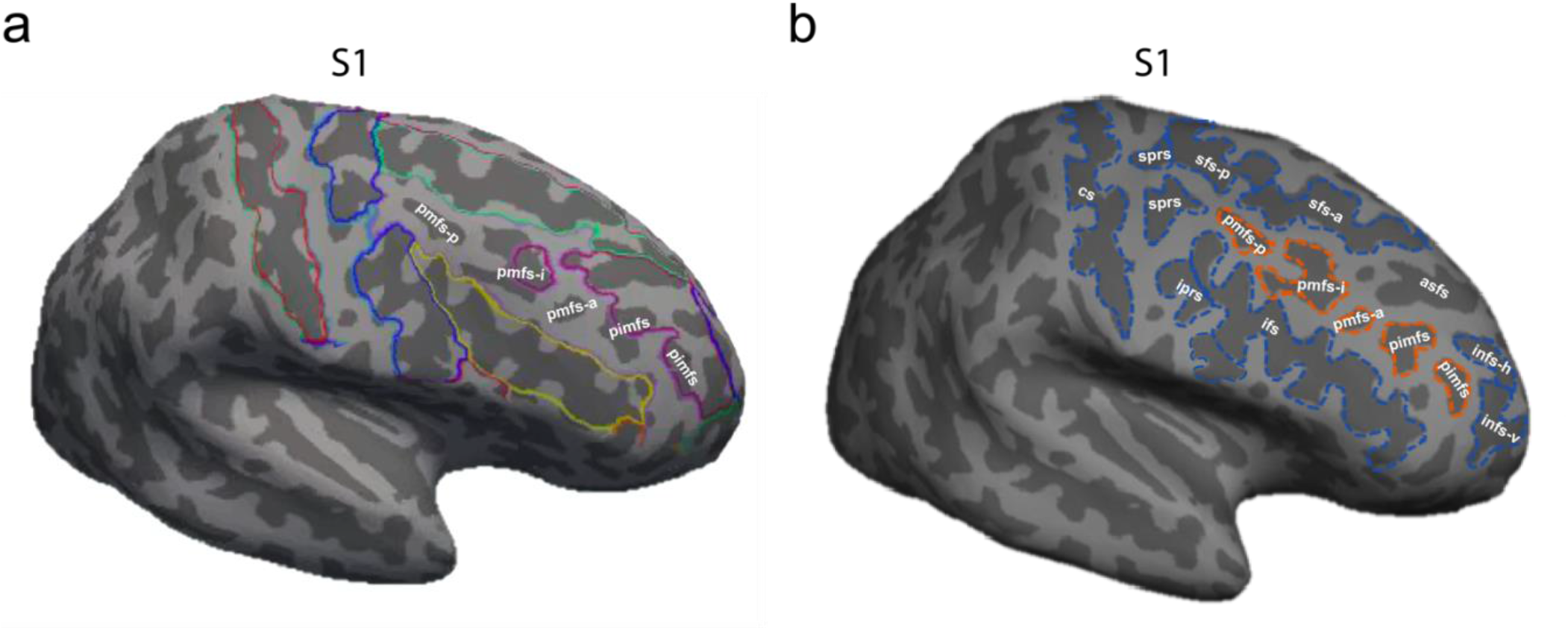
LPFC tertiary sulci are often omitted in commonly used atlases. **a.** Example inflated cortical surface reconstruction of a right hemisphere. Colors indicate sulcal and gyral definitions provided by automated methods^93,105^. The omitted tertiary sulci explored in the present study are labeled in white. While the automated
approach is useful for many studies, we manually defined sulci for our study as present approaches do not yet include tertiary sulci (labeled in white: *pmfs-p, pmfs-i, pmfs-a, pimfs*), and automated methods often include gyral components in the sulcal definitions. Red: central sulcus. Blue: pre-central sulcus. Yellow: inferior frontal sulcus. Turquoise: superior frontal sulcus. Magenta: fronto-marginal sulcus (which includes the horizontal and ventral components of the intermediate frontal sulcus, as well as portions of the pmfs-i, and what Petrides^14^ refers to as the *accessory superior frontal sulcus* (*asfs*; not examined in the present study). **b.** Example of manual sulcal definitions in the same subject as in **a.** Manual definitions capture both tertiary (*orange*) and primary (*blue*) sulci.

**Supplementary Table 1.** Variability in the number of pimfs components across individuals. Participants had 0, 1, or 2 pimfs components in each hemisphere. As a majority of participants in both samples had at least one *pimfs* component, our inclusion criteria was to include participants who had at least one *pimfs* component in each hemisphere (*Discovery*: 28/33, *Replication*: 27/28), which assures that all repeated measures statistics are balanced for effects of sulcus and hemisphere.

| Components | Discovery (n = 33) |  | Replication (n = 28) |  |
| --- | --- | --- | --- | --- |
|  | *left* | *right* | *left* | *right* |
| **0** | 2/33 | 3/33 | 1/28 | 0/28 |
| **1** | 15/33 | 18/33 | 3/28 | 9/28 |
| **2** | 16/33 | 12/33 | 24/28 | 19/28 |

**Supplementary Fig. 3.**
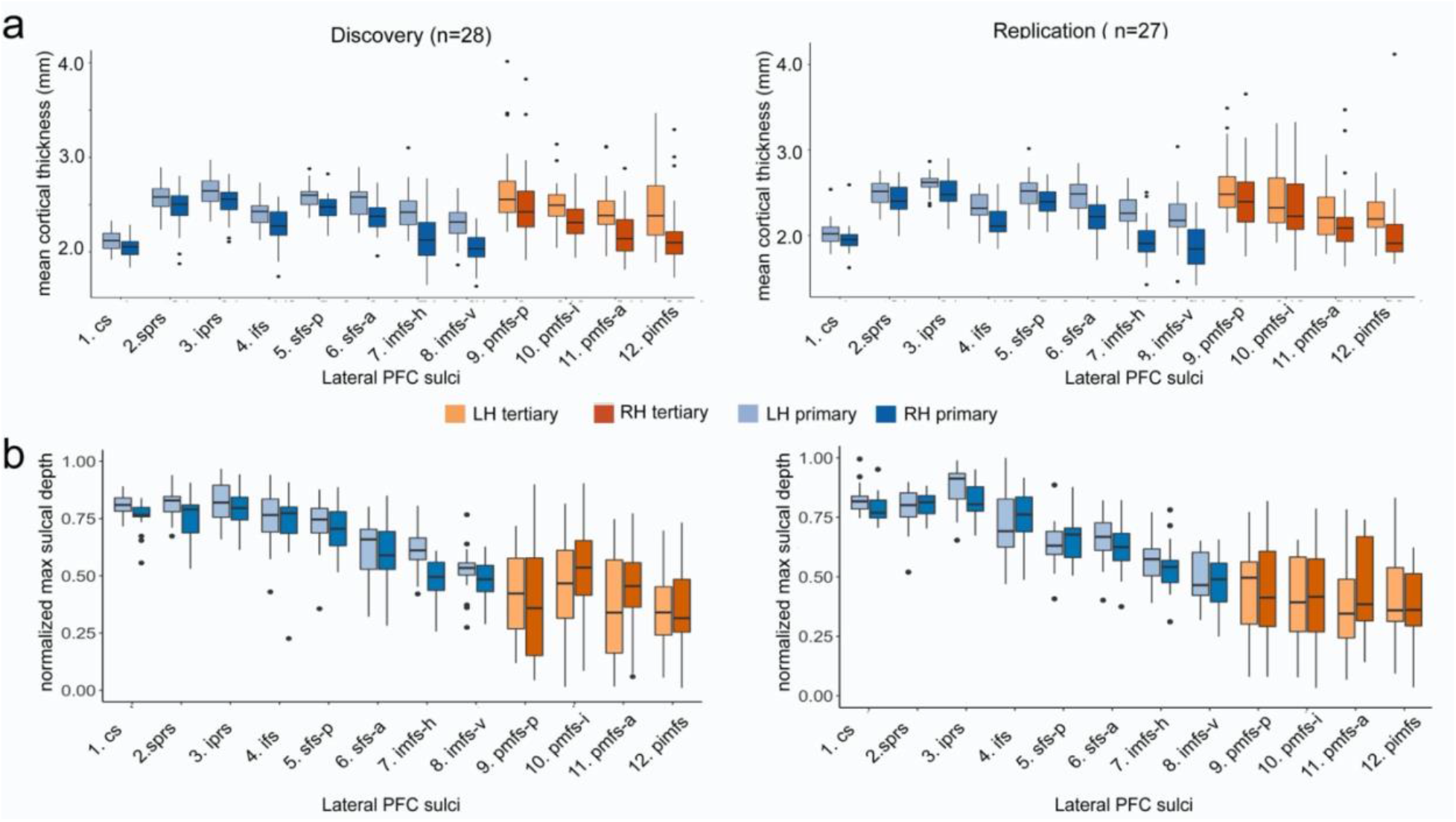
No difference in cortical thickness between tertiary and primary sulci in LPFC. **a.** *Left:* Mean cortical thickness for each of the 12 LPFC sulci in the *Discovery* sample. *Right:* Mean cortical thickness for each of the 12 LPFC sulci in the *Replication* sample. Tertiary sulci (*orange*) and primary sulci (*blue*) do not significantly differ in cortical thickness in either sample. **b.** *Left:* Maximum sulcal depth for each of the 12 LPFC sulci in the *Discovery* sample. *Right:* Maximum sulcal depth for each of the 12 LPFC sulci in the *Replication* sample. Depth was normalized by the deepest point in the individual’s hemisphere.

**Supplementary Fig. 4.**
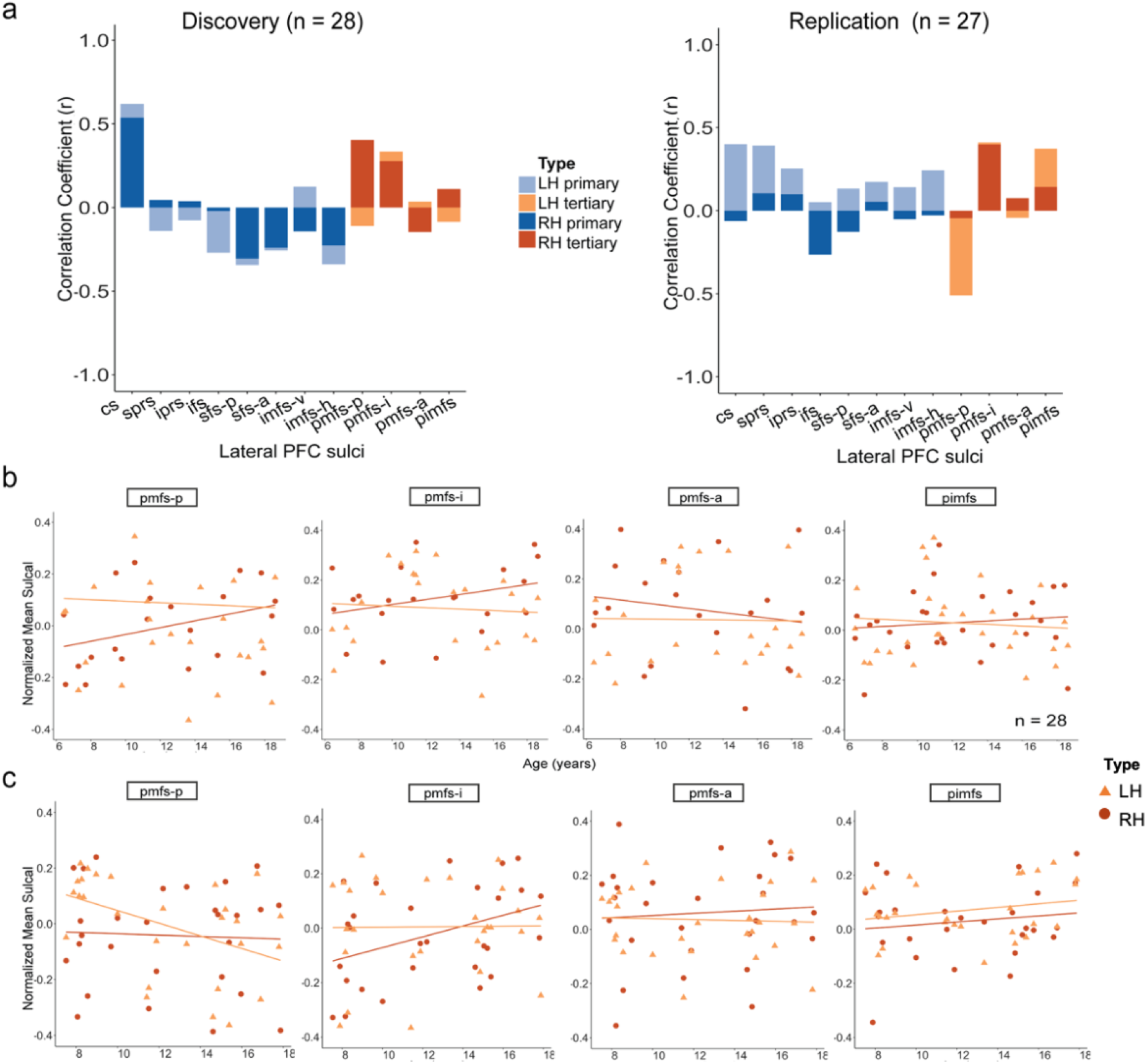
Morphological and behavioral associations with age in both Discovery and Replication samples. **a.** Correlation between age and sulcal depth in the *Discovery* (*left*) and *Replication* (*right*) samples. Each bar represents the correlation coefficient (Pearson’s *r*) between sulcal depth and age for each sulcus (*orange:* tertiary; *blue:* primary) in the left (lighter shades) and right (darker shades) hemispheres. There is not a clear relationship between sulcal depth and age that is generalizable among LPFC sulci. **b-c.** Scatterplots showing the association between age and sulcal depth for each of the 4 tertiary sulci explored in the present study in each hemisphere (left: lighter shades; right: darker shades) for individual participants in the *Discovery* (**b**) and *Replication* (**c**) samples. Age does not account well for individual variability in LPFC tertiary sulcal depth.

**Supplementary Fig. 5.**
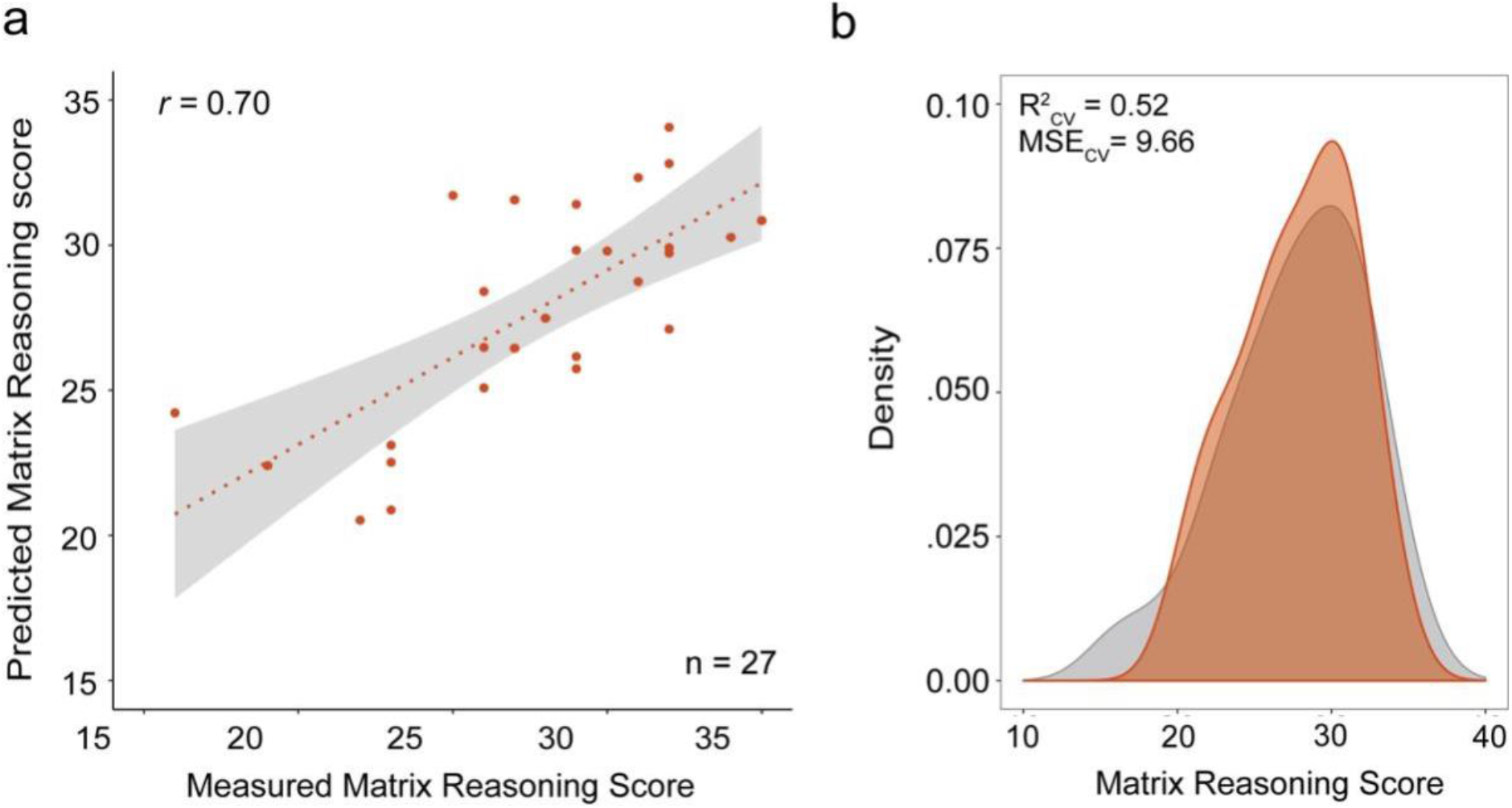
Predicted matrix reasoning score in the Replication sample from three tertiary sulci (pmfs-i, pmfs-a, pimfs). **a.** Spearman’s correlation between measured and predicted Matrix Reasoning scores in the *Replication* sample for the model including all three tertiary sulci identified in the *Discovery* sample (*pmfs-i_RH_, pmfs-a_RH_, pimfs_RH_*).**b**. Density plot showing model fit. *orange*: The distribution of predicted scores from this model. *gray:* the distribution of measured Matrix Reasoning scores.

**Supplementary Fig. 6.**
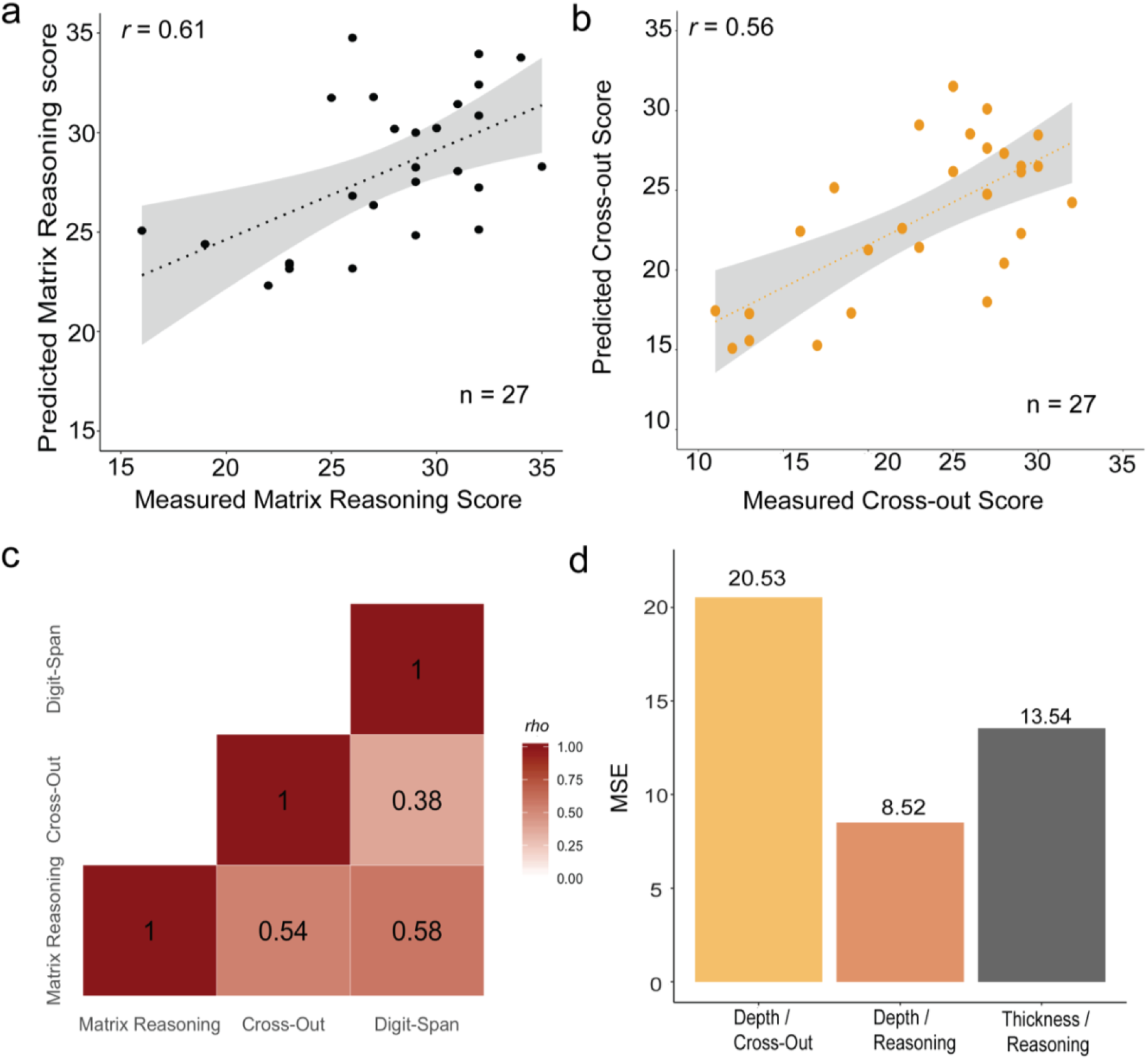
Tertiary sulcal depth more strongly relates to reasoning than cortical thickness and this relationship shows behavioral preference over other cognitive measures. **a.** Thickness was used in place of depth to predict Matrix Reasoning in the *Replication* sample. The model was fit with looCV. Spearman’s correlation between measured and predicted Matrix Reasoning scores in the *Replication* sample for the best model (pmfs-i_RH_ + pimfs_RH_ + age). b. The same depth model was used to predict Cross-Out score instead of Matrix Reasoning in the *Replication* sample. Spearman’s correlation between measured and predicted Cross-Out scores in the *Replication* sample using the best performing depth model (pmfs-i_RH_ + pimfs_RH_ + age). c. Correlation (Spearman’s *rho*) between Matrix Reasoning, Cross-out (Processing speed), and Digit span (Working memory). Digit span was included as an additional comparison measure in the Replication sample. Digit span was not predicted by the depth model (and was not explored further (R^2^_cv_ = 0.10)). d. MSE_cv_ for the thickness and Cross-Out score models compared to the analogous depth model. Tertiary sulcal depth offered substantially better predictions than cortical thickness. The depth model did not generalize to processing speed.

## Acknowledgments

This research was supported by a T32 HWNI training grant and an NSF-GRFP fellowship (Voorhies), as well as start-up funds from UC Berkeley (Weiner). Funding for the original data collection and curation was provided by NINDS R01 NS057156 (Bunge, Ferrer) and NSF BCS1558585 (Bunge, Wendelken). Data analysis was supported by NICHD R21HD100858 (Weiner, Bunge). We thank Ishana Raghuram for assistance with sulcal labeling and former members of the Bunge laboratory for assistance with data collection, and the families who participated in the study.

